# PspA adopts an ESCRT-III-like fold and remodels bacterial membranes

**DOI:** 10.1101/2020.09.23.309765

**Authors:** Benedikt Junglas, Stefan T. Huber, Thomas Heidler, Lukas Schlösser, Daniel Mann, Raoul Hennig, Mairi Clarke, Nadja Hellmann, Dirk Schneider, Carsten Sachse

## Abstract

PspA is the main effector of the phage shock protein (Psp) system and preserves the bacterial inner membrane integrity and function. Here, we present the 3.6 Å resolution cryo-EM structure of PspA assembled in helical rods. PspA monomers adopt a canonical ESCRT-III fold in an extended open conformation. PspA rods are capable of enclosing lipids and generate positive membrane curvature. Using cryo-EM we visualized how PspA remodels membrane vesicles into μm-sized structures and how it mediates the formation of internalized vesicular structures. Hot spots of these activities are zones derived from PspA assemblies, serving as lipid transfer platforms and linking previously separated lipid structures. These membrane fusion and fission activities are in line with the described functional properties of bacterial PspA/IM30/LiaH proteins. Our structural and functional analyses reveal that bacterial PspA belongs to the evolutionary ancestry of ESCRT-III proteins involved in membrane remodeling.

## Introduction

Membrane remodeling and re-shaping is crucial for cell viability in all domains of life. In order to maintain the integrity of internal membranes under conditions such as heat stress, osmotic stress, organic solvents or phage infections, many bacteria activate the phage shock protein (Psp) system (Brissette et al., 1990; Kobayashi et al., 1998; Weiner et al., 1991). The currently best studied *E. coli* Psp*-*system consists of seven genes: The *pspABCDE* operon, the regulation factor *pspF* and the effector *pspG* (Brissette et al., 1991; Elderkin et al., 2002; Lloyd et al., 2004). The Psp response is triggered during dissipation of the proton motive force and secretin mislocation (Darwin, 2005; Engl et al., 2011; Joly et al., 2010), which both indicate defects in the structure and integrity of the bacterial cytoplasmic membrane. PspA is the key effector of the Psp system and in *E. coli* interacts with the cytoplasmic membrane sensor proteins PspB and PspC (Darwin, 2005; Joly et al., 2010). Free PspA can bind peripherally to bacterial inner membranes by interacting with negatively charged membrane lipids (Jovanovic et al., 2014) and thereby counteracts membrane stress (Joly et al., 2010; Manganelli and Gennaro, 2017). This way, membrane-bound PspA can, for instance, block proton leakage of isolated damaged membranes (Kobayashi et al., 2007). In support, members of the PspA family are thought to tether and seal stress-induced membrane lesions, presumably by triggering lipid bilayer fusion (Siebenaller et al., 2020). Indeed, a membrane fusion activity has been reported previously for the PspA homolog IM30 (inner membrane associated protein of 30 kDa, alternatively known as Vipp1) of the cyanobacterium *Synechocystis* sp. PCC 6803 (Hennig et al., 2015). More recently, a membrane protecting protein carpet mechanism has been proposed for IM30 (Junglas et al., 2020).

While the Psp response system appears to be conserved in Gram-negative bacteria, Psp homologs have also been identified in cyanobacteria, archaea as well as in chloroplasts (Manganelli and Gennaro, 2017). Yet, solely homologs of PspA appear to be strictly conserved and thus critical for membrane protection and remodeling (Manganelli and Gennaro, 2017). The homologous proteins PspA, IM30/Vipp1 (chloroplasts and cyanobacteria) and LiaH (*Firmicutes, Bacillus and Listeria*) are part of a bacterial membrane remodeling family that shares many structural and functional features for maintaining membrane integrity (Thurotte et al., 2017; Vothknecht et al., 2012). PspA/LiaH/IM30 proteins form large ring and rod-like homo-oligomeric structures (>1 MDa), and share predicted secondary structures and conserved sequence motifs (Bultema et al., 2010; Fuhrmann et al., 2009; Hankamer et al., 2004; Male et al., 2014; Saur et al., 2017; Theis et al., 2019, 2020; Thurotte et al., 2017; Vothknecht et al., 2012; Wolf et al., 2010). While the structure of an extended α-helical hairpin formed by the 2^nd^ and 3^rd^ *Eco*PspA helices was resolved by X-ray crystallography (Osadnik et al., 2015), the full-length PspA structure is still unknown.

Although most membrane-remodeling proteins such as epsin, dynamin or BAR domain-containing proteins were first identified in eukaryotes (Ford et al., 2002; Frost et al., 2009; Praefcke and McMahon, 2004), membrane remodeling is a common process in all three domains of life. For instance, e*ndosomal sorting complexes required for transport* (ESCRT) were originally characterized in eukaryotic membrane trafficking events of multivesicular body biogenesis (Katzmann et al., 2001), viral budding (Garrus et al., 2001) and cytokinesis (Carlton and Martin-Serrano, 2007) with the common topology of moving membrane away from the cytosol. More recently, they have been shown to be involved in a series of diverse membrane remodeling processes in eukaryotic cells, which include nuclear envelope sealing (Vietri et al., 2015), plasma membrane repair, lysosomal protein degradation (Zhu et al., 2017) and closure of autophagosomes (Takahashi et al., 2018; Zhen et al., 2020). Despite this functional diversity, the responsible core proteins belong to a single conserved group of ESCRT-III proteins. ESCRT-III proteins comprise approx. 220 amino acids and share seven predicted α-helices (helices α0 - α6). Structural analyses of truncated CHMP3 revealed a compact structure made of a four-helix bundle comprising of a long α-helical hairpin motif (Muzioł et al., 2006). More recently, polymers assembled from full-length ESCRT-III were elucidated at near-atomic resolution using cryo-EM: human IST1 and CHMP1B form a hetero-polymer with a compact and extended conformation forming a tubular assembly (McCullough et al., 2015), whereas yeast Vps24 assembles in a double-stranded filament composed of domain-swapped dimers in the extended conformation (Huber et al., 2020). The structures of IST1/CHMP1B tubes confirmed the presence of a lipid bilayer in the lumen of the assemblies (Nguyen et al., 2020). Thus, IST1/CHMP1B is capable of constricting vesicles and inducing positive membrane curvature suitable for opposite ESCRT topology membrane processes. It has been shown that mixing of different ESCRT proteins in a particular order is required to assemble hetero-polymeric complexes that drive membrane remodeling *in vitro* (Huber et al., 2020; Pfitzner et al., 2020). Downstream of the ESCRT-III activity is the AAA-ATPase Vps4 that promotes assembly and disassembly of ESCRT-III complexes and thereby contributes to membrane remodeling (Caillat et al., 2019).

Despite the known critical roles of ESCRT-III proteins in the Archaea and Eukarya kingdom, members of a conserved bacterial ESCRT family have not yet been identified, neither by sequence analysis, structural comparison nor by functional characterization. In the current study, we determined the 3.6 Å resolution cryo-EM structure of PspA assembled in helical tubes. The built atomic model reveals that PspA adopts a canonical ESCRT-III fold, making up the wall of the tubular assemblies. When PspA is reconstituted with small lipid vesicles, it induces vesicle growth and formation of internalized membrane structures, indicating membrane fusion as well as fission activities. The cryo-EM structure combined with biochemical lipid remodeling experiments show that bacterial PspA is a *bona fide* member of an evolutionary conserved ESCRT-III family.

## Results

### PspA rods are tubular assemblies with helical symmetry

In order to structurally investigate the membrane protective mechanism of PspA, we overexpressed PspA of the cyanobacterium *Synechocystis* sp. PCC 6803. Expression of a fluorescent *Synechocystis* PspA variant in a Δ*pspA E. coli* strain resulted in the formation of multiple, large PspA assemblies *in vivo*, which were preferably localized at the cell poles **(Figure S1A**), supporting previous observations of large inner membrane associated PspA oligomers (Engl et al., 2009; Jovanovic et al., 2014; Yamaguchi et al., 2013). Subsequently, we visualized purified PspA using cryo-EM. The acquired micrographs show elongated rod assemblies with around 20 nm width and varying lengths of up to several hundreds of nanometers (**Figure 1A**). Closer analysis of the 2D class average from a homogeneous subset (**Figure S1B&C)** revealed a basic Christmas-tree pattern with a longitudinal repeat every 30 Å indicative of a repeating pitch typical for helical assemblies (**Figure 1B**). The corresponding power spectrum showed a layer line pattern diffracting to approx. 7 Å resolution and revealed the helical organization with a helical rise and rotation of 2.9 Å and 35.3º per subunit (**Figure 1C, Figure S1D**). Using a total of 139,300 asymmetric units, we then determined the cryo-EM structure at an overall resolution of 3.6 Å (**Figure 1D**), with resolution variation between 3.5 Å and 4.7 Å at 50 Å and 100 Å radial distance from the helical axis, respectively (**Figure 1E**). After density segmentation, we found that a single subunit has an elongated shape of 180 × 20 Å dimension, which is positioned perpendicularly to the rod axis (**Figure 1F**). When seen in top view, the subunit extends over one third of the rod’s cross section. In side view, the subunits are arranged like overlapping bricks that constitute the wall of the helical rod (**Figure 1G**).

**Figure 1:**
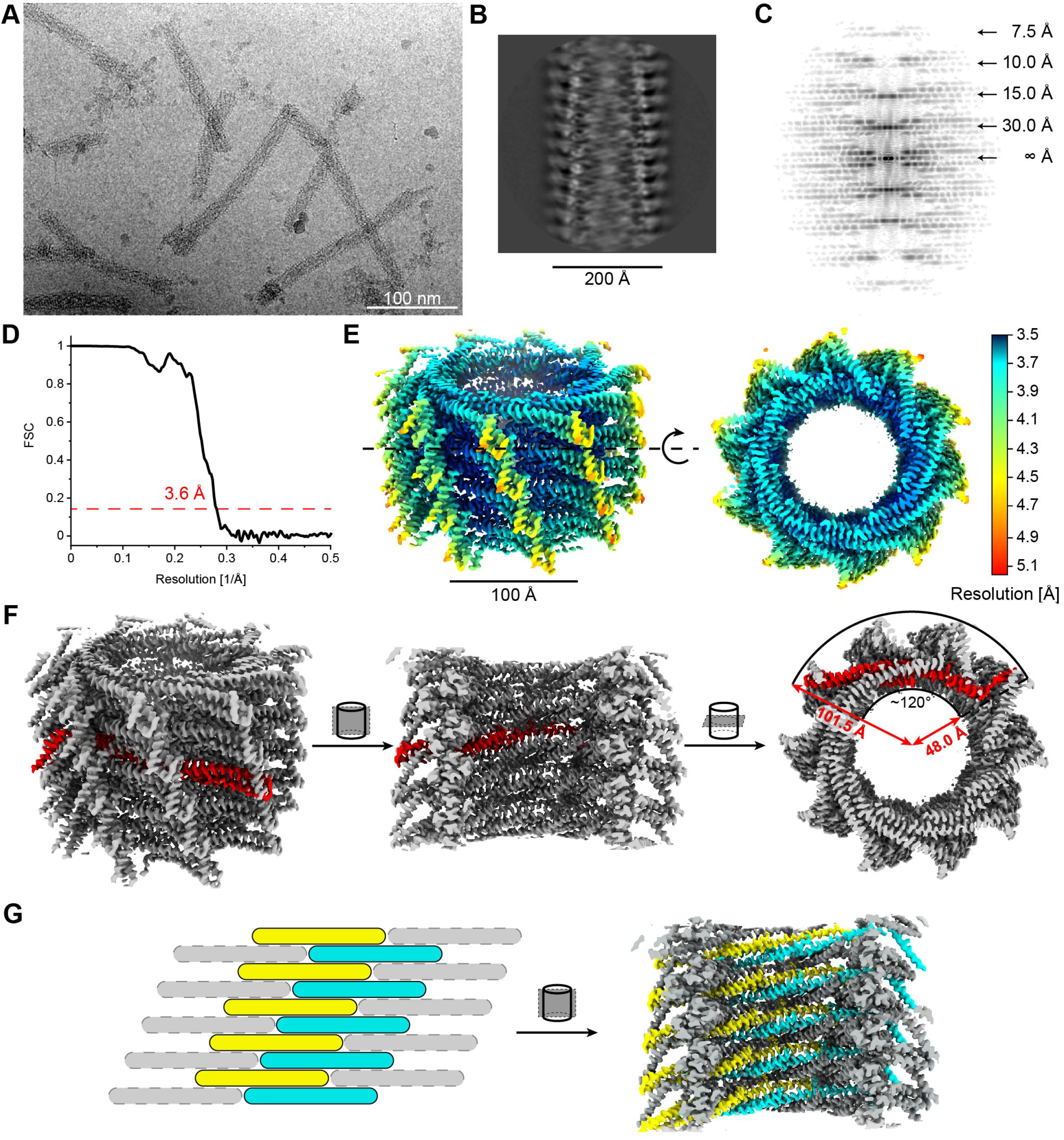
Cryo-EM structure of helical PspA assemblies. (A) Representative electron cryo-micrograph of PspA rods. (B) Detailed class average of Christmas-tree appearance and (C) corresponding power spectrum indicating layer lines up to 7 Å resolution. (D) Fourier shell correlation supporting 3.6 Å resolution at the 0.143 cutoff. (E) Local resolution estimates are mapped onto the surface of the PspA cryo-EM structure. The resolution varies from 3.5 Å at the inner wall to 4.7 Å at the C-terminal helix (see also **Table 1** and **Figure S1B–D**). (F) Left: side view of cryo-EM density with a single segmented subunit in red. Center: open cylinder side view. Right: top view. (G) Left: schematic view of subunits stacked like bricks forming a wall. Right: corresponding open half-cylinder view of stacked subunits forming the cylinder wall.

**Table 1.**
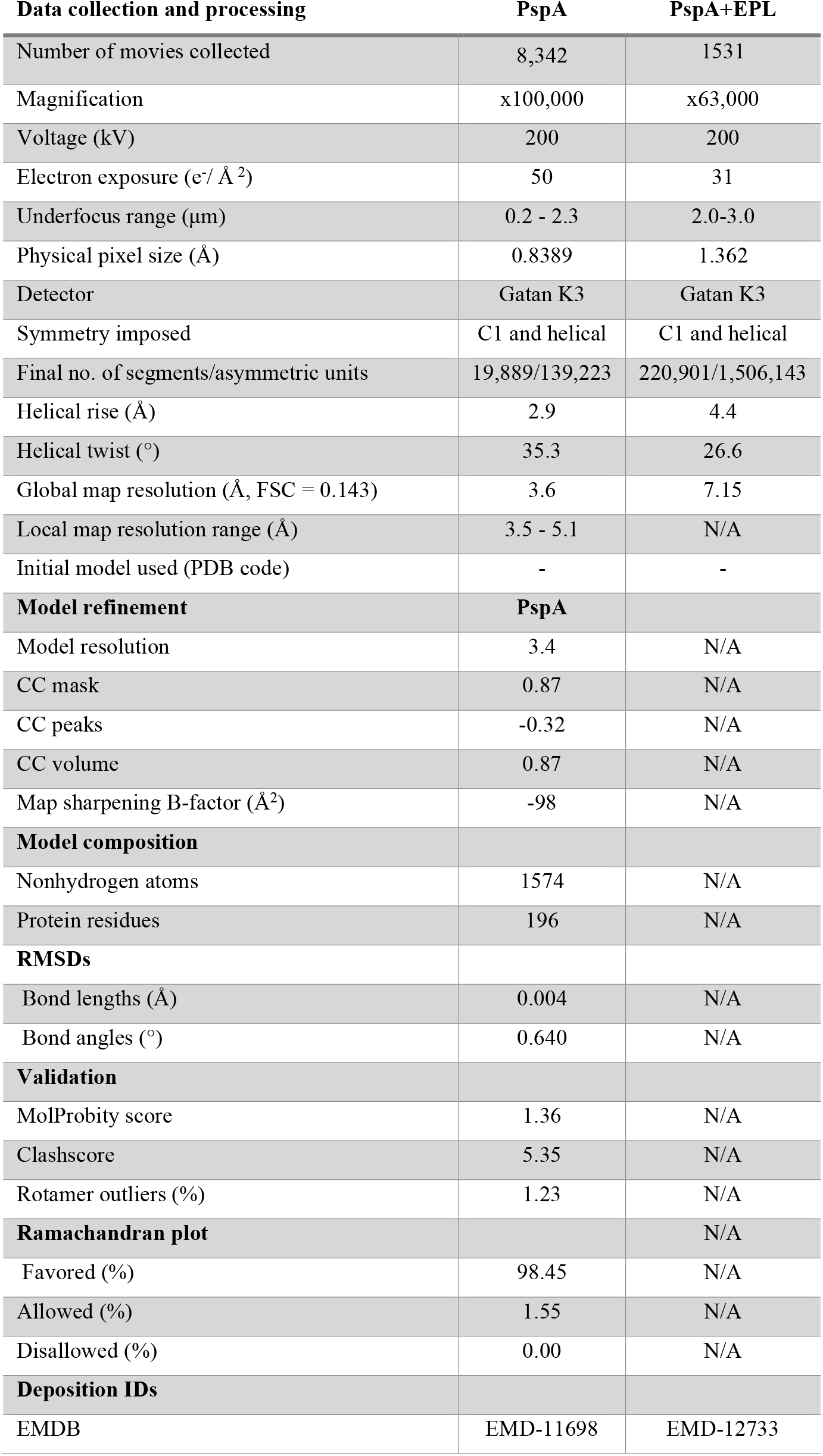
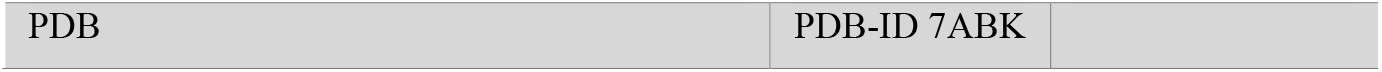
PspA cryo-EM structure determination.

| **Data collection and processing** | **PspA** | **PspA+EPL** |
| --- | --- | --- |
| Number of movies collected | 8,342 | 1531 |
| Magnification | x100,000 | x63,000 |
| Voltage (kV) | 200 | 200 |
| Electron exposure (e<sup>-</sup>/ Å<sup>2</sup>) | 50 | 31 |
| Underfocus range (μm) | 0.2 - 2.3 | 2.0-3.0 |
| Physical pixel size (Å) | 0.8389 | 1.362 |
| Detector | Gatan K3 | Gatan K3 |
| Symmetry imposed | C1 and helical | C1 and helical |
| Final no. of segments/asymmetric units | 19,889/139,223 | 220,901/1,506,143 |
| Helical rise (Å) | 2.9 | 4.4 |
| Helical twist (°) | 35.3 | 26.6 |
| Global map resolution (Å, FSC = 0.143) | 3.6 | 7.15 |
| Local map resolution range (Å) | 3.5 - 5.1 | N/A |
| Initial model used (PDB code) | - | - |
| **Model refinement** | **PspA** |  |
| Model resolution | 3.4 | N/A |
| CC mask | 0.87 | N/A |
| CC peaks | -0.32 | N/A |
| CC volume | 0.87 | N/A |
| Map sharpening B-factor (Å<sup>2</sup>) | -98 | N/A |
| **Model composition** |  |  |
| Nonhydrogen atoms | 1574 | N/A |
| Protein residues | 196 | N/A |
| **RMSDs** |  |  |
| Bond lengths (Å) | 0.004 | N/A |
| Bond angles (°) | 0.640 | N/A |
| **Validation** |  |  |
| MolProbity score | 1.36 | N/A |
| Clashscore | 5.35 | N/A |
| Rotamer outliers (%) | 1.23 | N/A |
| **Ramachandran plot** |  | N/A |
| Favored (%) | 98.45 | N/A |
| Allowed (%) | 1.55 | N/A |
| Disallowed (%) | 0.00 | N/A |
| **Deposition IDs** |  |  |
| EMDB | EMD-11698 | EMD-12733 |

### PspA adopts an elongated four-helix core with an accessory C-terminal helix

Using the segmented density, we built an atomic model of a PspA monomer, covering residues 22 to 217 arranged in five α-helices connected by loops (**Figure 2A–C**). At one end of the elongated subunit, we identified a density that corresponds to the long α-helical hairpin PspA fragment (PDB-ID 4WHE, (Osadnik et al., 2015)), independently confirming the chosen handedness of the cryo-EM map (**Figure 2D**). The best resolved density accommodates helices α1-α4 (25-185) (**Figure 2E, F**). We did not find density corresponding to a predicted N-terminal helix α0 (1-21) (**Figure S1E**), yet, we accommodated the amphipathic helix α5 (191-215) based on the weaker density attached to the outer wall of the helical rod (**Figure 2G, Figure S1F**). Together, helices α1-α4 constitute the elongated four-helix core of the PspA molecules that form the wall of the tubular rod structure via a brick-like stacking. A total of 10.19 subunits are wound in a right-handed manner to form a single 360º turn with a pitch of 30 Å (**Figure 2H**). The top view of the assembly reveals that the PspA core helices are arranged like the diaphragm of an iris aperture pointing to the outside of the rod (**Figure 2I**) while the amphipathic paddle helix α5 at the C-terminus emanates from the wall to the outside of the helical rod (**Figure 2J**). Interestingly, at both ends of the PspA core, first at the tip of the hairpin between helix α1 and α2 (70-90) and second at the opposite end of helix α4 (167-180), we find highly conserved (>90 % identity) residues among the PspA/IM30 family of proteins (**Figure 2K&L, Figure S1G**) suggesting an evolutionary importance of these intersubunit stabilizing residues. Previously, mutation of these residues abolished formation of *Syn*IM30 rods, a closely related PspA homolog (Hennig et al., 2015; Saur et al., 2017). Therefore, we mutated the hydrophobic residues A75 and A78 and analyzed the PspA_A75S/A78S preparations using negative staining EM. While PspA_A75S/A78S is still able to form rod structures, our image data suggest that mutant rods are less stable, in particular after refolding in the presence of 75 mM NaSCN when the mutant does not form rods anymore (**Figure S2A&B**). Moreover, we computed the Coulomb potential of the atomic model PspA assembly and found that both ends show opposite charges with a basic and acidic surface, respectively (**Figure 2M**), resulting in a polar assembly that has very different binding surfaces at each end of the assembly.

**Figure 2:**
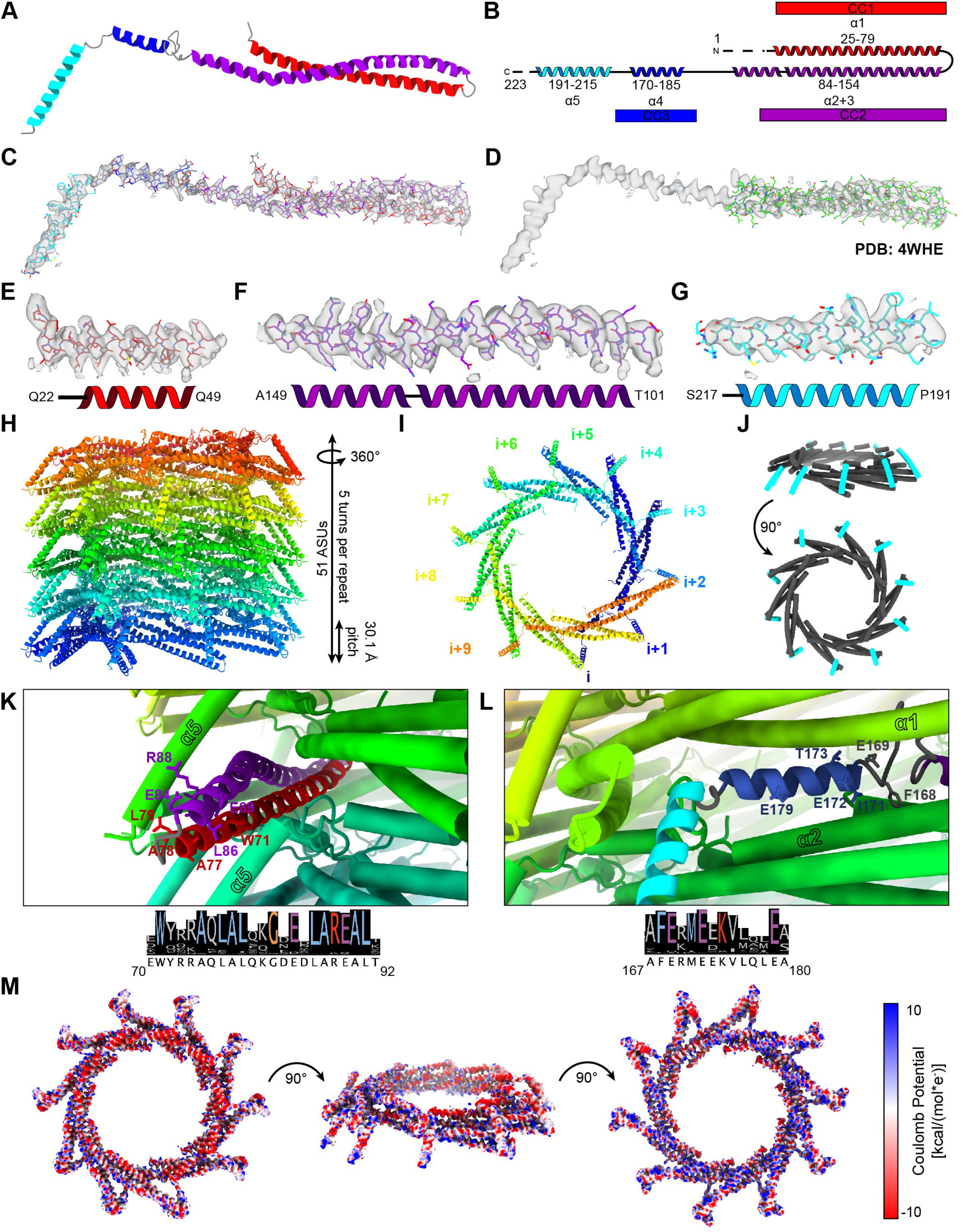
Atomic model and architecture of the helical PspA assembly. (A) Atomic model of PspA (22-217) with colored α-helices in ribbon presentation (α1: red, α2/α3: purple, α4: blue, α5: cyan; see also **Table 1**). (B) Secondary structure overlaid in the same color scheme as in (A) (see also **Figure S1G**) (Helix nomenclature follows the ESCRT-III convention). (C) Segmented cryo-EM density superimposed on built atomic model of PspA. (D) Rigid body fit of the *Eco*PspA (25-141) X-ray structure PDB-ID 4WHE. (E) Density of helix α1, (F) helix α2/α3 and (G) helix α5 at lower resolution. (H) The rod assembly in side view: elongated monomers (in rainbow colors) assemble to form a right-handed helix of 10.19 subunits per turn, *i.e.* corresponding to a pitch of 30.1 Å. (I) Top view of the assembly corresponding to one turn of stacked monomers i, i+1, i+2 … i+9 (in rainbow colors). (J) Helix α5 (cyan) forms a paddle that emanates from the wall to the outside of the helical rod. (K) Scene 1 (residues 70-92) and (L) Scene 2 (residues 167-180) show the structural context of highly conserved side chains (color code as in B) in the PspA assembly (color code as in H). (M) Electrostatic surface potential of PspA rods. The assembly consists of two unique ends with basic and acidic surfaces.

### PspA shares the basic domain architecture with ESCRT-III proteins

Based on the here determined cryo-EM structure of PspA, we found one particular structural feature, *i.e.* the helix α1/α2 hairpin packing against a perpendicular helix α5, reminiscent of previously resolved structures of eukaryotic membrane-remodeling ESCRT-III proteins. Therefore, we closer examined their apparent similarity by sequence alignment and compared cyanobacterial PspA and IM30 proteins with yeast Vps24 as well as human IST1 and CHMP1B (**Figure 3**), for which high-resolution cryo-EM structures have been determined. Interestingly, there are several stretches that show high sequence similarities (Quality score >4, **Figure S2C**). Furthermore, structural superposition of the α1/α2 hairpin region yields CA RMSDs with *Eco*PspA (PDB-ID 4WHE, (Osadnik et al., 2015)), IST1 (PDB-ID 6TZA, (Nguyen et al., 2020)), CHMP1B (PDB-ID 6TZ9, (Nguyen et al., 2020)) and Vps24 (PDB-ID 6ZH3, (Huber et al., 2020)) reporting average RMSD values of 1.6 Å, 2.3 Å, 2.2 Å and 1.8 Å, respectively (**Figure S2D**). CHMP1B, Vps24 and PspA adopt elongated conformations, whereas IST1’s structure was determined as a compact fold (**Figure 4A**). The corresponding resolved higher-order structures show that all of their assemblies remain unique (**Figure 4B**) while the fundamental organization of the α-helices is conserved between all ESCRT-III proteins with helix-to-helix loops variations. Nevertheless, common to all structures is that the C-terminal helix α5 packs approx. perpendicularly against the α1/α2 hairpin motif (**Figure 4C**). Based on the identified structural similarities between the here determined bacterial PspA cryo-EM structure and members of the eukaryotic ESCRT-III family, we conclude, PspA may have shared evolutionary ancestry with ESCRT-III proteins.

**Figure 3:**
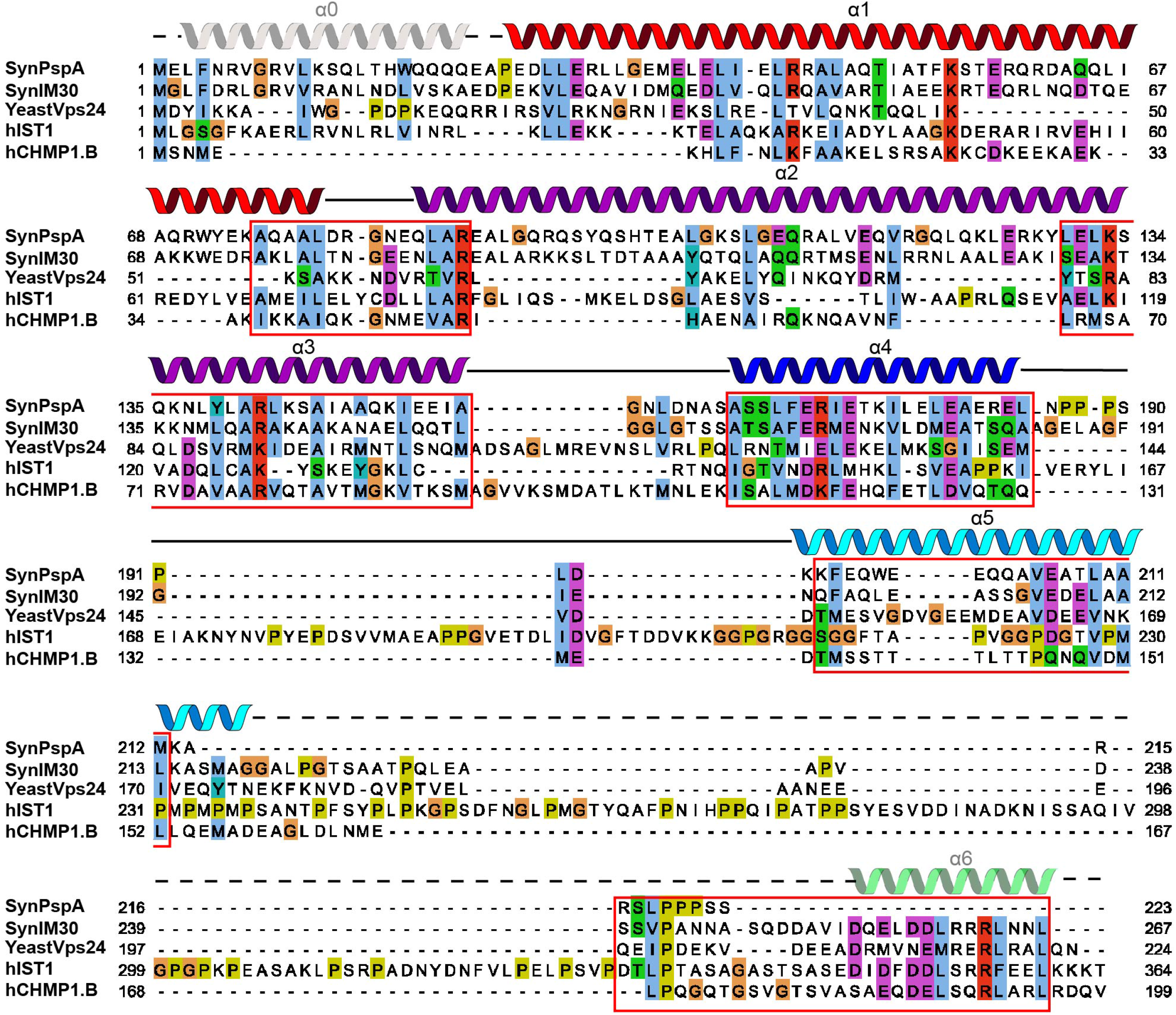
Sequence comparison of PspA/IM30 with ESCRT-III proteins. Sequence alignment of *Syn*PspA, *Syn*IM30, Yeast Vps24, human IST1 and human CHMP1B superimposed with the secondary structures denoted by ESCRT-III nomenclature. Residues with similarity >30 % are colored according to their properties (ClustalX color scheme). Stretches of high similarity are highlighted with red boxes. Alignment scores for the boxed areas are shown in **Figure S2C**. Sequences were aligned using the T-Coffee webserver (Di Tommaso et al., 2011) and visualized with JalView (Waterhouse et al., 2009).

**Figure 4:**
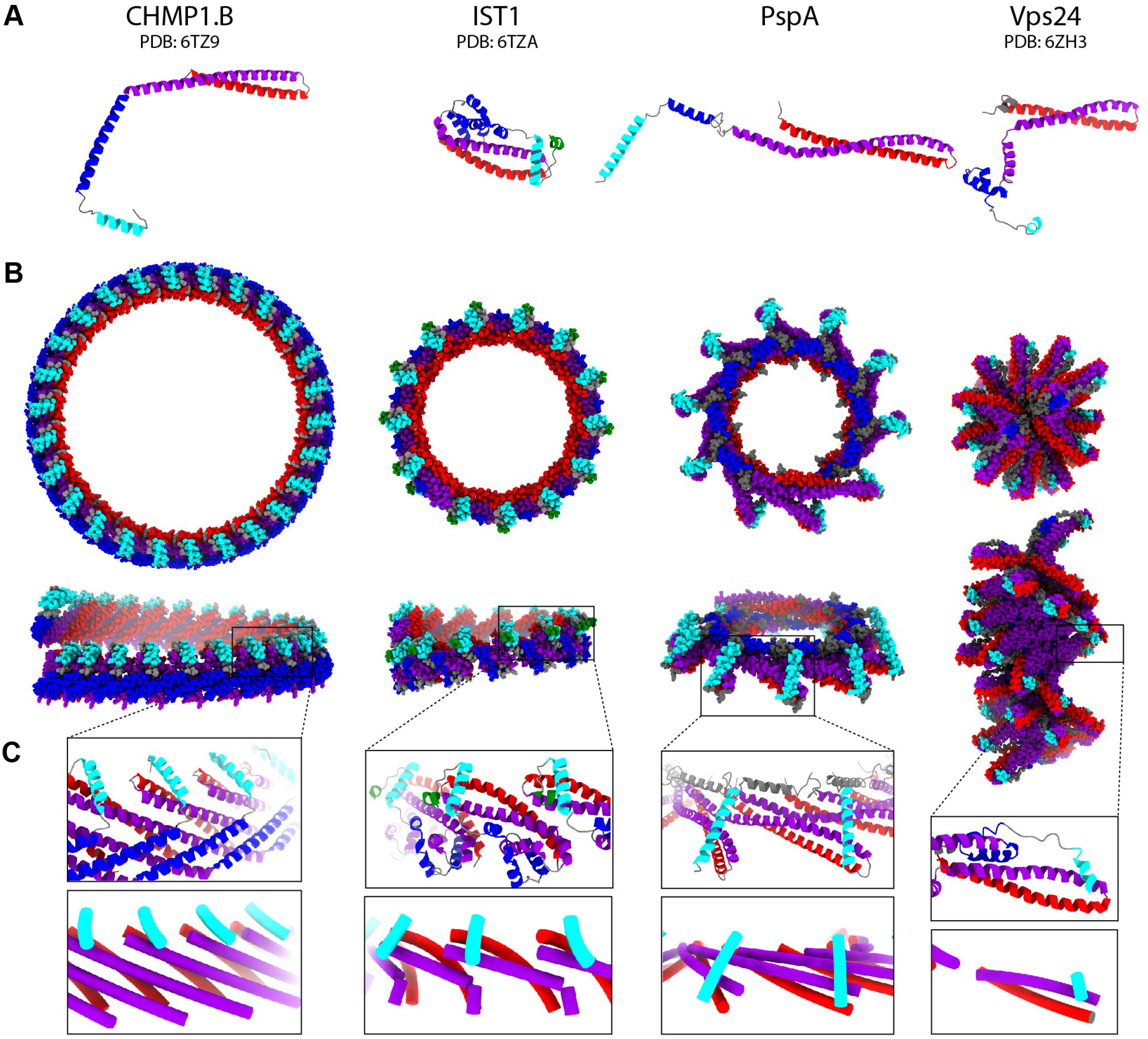
Comparison of PspA and ESCRT-III structures. (A) Views of CHMP1B, IST1, PspA and Vps24 monomers with colored α-helices in ribbon presentation (α1: red, α2/α3: purple, α4: blue, α5: cyan). (B) Helical assemblies of ESCRT-III proteins and PspA in sphere representation (color code as in A). Upper row: top views, lower row: side views. (C) Details of the ESCRT-III and PspA assemblies showing a conserved motif with helix α5 being packed approx. perpendicularly against the α1/α2 hairpin motif.

### PspA binds to lipid bilayers and converts small vesicles into large membrane structures

Due to the observed structural homology of PspA with ESCRT-III proteins, we tested whether PspA has a membrane remodeling activity in analogy to the ESCRT proteins. Therefore, we incubated preformed PspA rods with small unilamellar (SUVs) vesicles prepared from *E. coli* polar lipid extract (EPL) and analyzed the mixture using cryo-EM. In the cryo-micrographs of the PspA/EPL mixtures, PspA rods in solution as well as a large fraction of vesicles with increased diameters of >2 μm were observed. The large vesicles often have round and elongated shapes, enclose smaller deformed vesicles and can even have more complex internalized lipid structures (**Figure 5A&B**, **Figure S3A**). As expected, in the SUV control we visualized a population of small vesicles with 20 - 200 nm diameter (**Figure 5C**). Vesicle dimension and perimeter measurements indicate a significant increase of membrane length upon addition of PspA rods (**Figure 5D–F**), suggesting the fusion of several smaller vesicles into one large vesicle. In many cases, the lipid bilayer appears to be interrupted by fuzzy stretches that have PspA protein density as well as PspA rods associated with the membrane (**Figure 5B**, **Figure S3B**). Furthermore, when analyzing the membrane fine structure of large vesicles by 2D classification, the peak-to-peak separation of the bilayer increased from 3.0 to 5.4 nm on average when compared with the lipid samples alone (**Figure S3C&D**). Interestingly, the increased bilayer leaflet separation is only found in vesicles with perimeters larger than 175 nm, suggesting a post-fusion membrane structure induced by PspA. Additional micrographs of large unilamellar vesicles (LUVs) (**Figure S4A**) or the LUV/PspA mixture confirm the observation of an increased leaflet separation of 5.7 nm for fused vesicles (**Figure S4B&C**). In fact, when we analyzed the interaction of PspA with model membranes using fluorescence spectroscopy, we observed binding of PspA to negatively charged PG-containing membranes (**Figure 5G**), as well as net uncharged PC membranes (**Figure S5A&B**). Destabilized rods (PspA_A75S/A78S) also bound to model membranes, even with higher apparent affinity, supporting the observation that PspA membrane binding involves rod disassembly.

**Figure 5:**
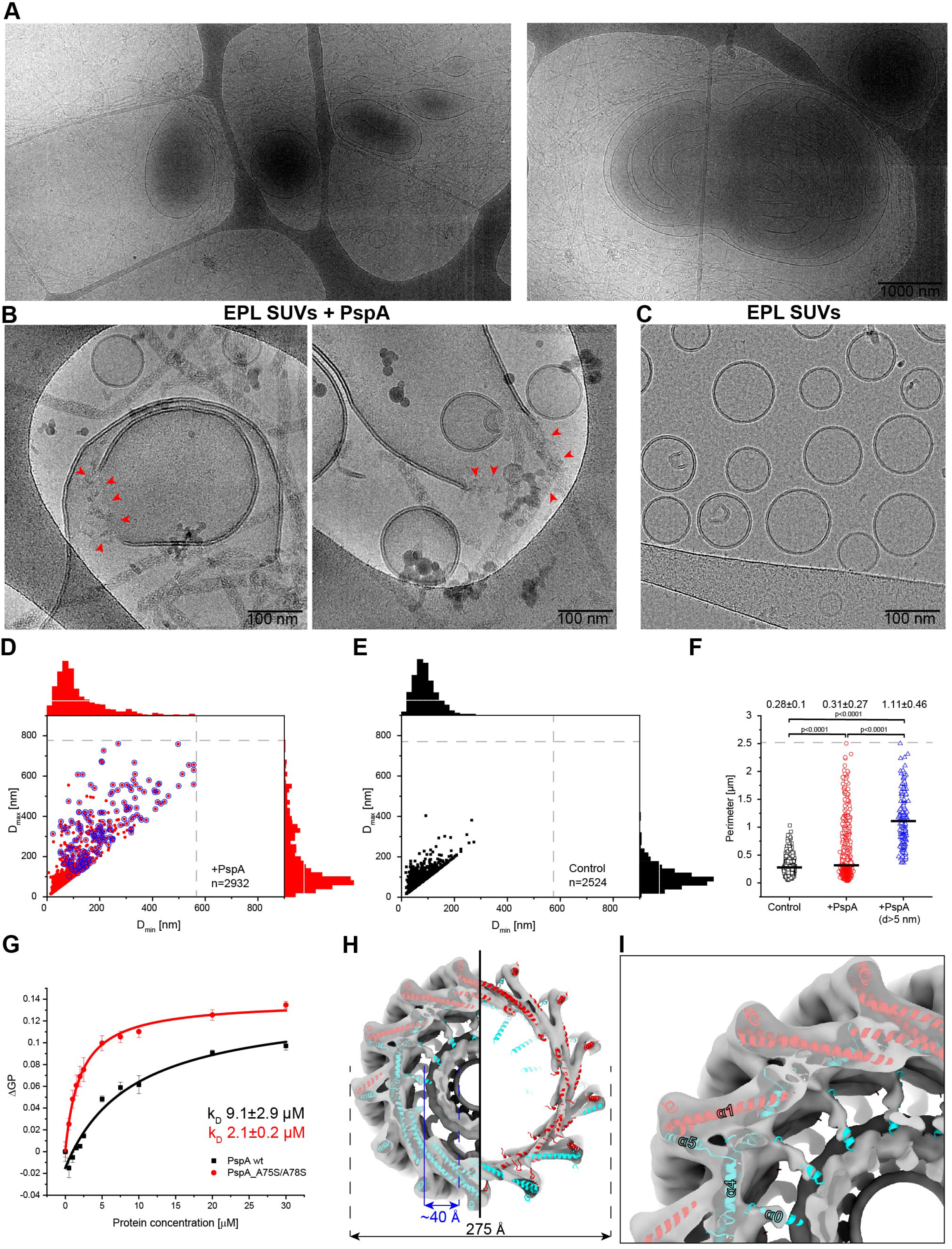
Membrane binding and remodeling by PspA. (A) Electron cryo-microscopy (cryo-EM) overview images of small unilamellar vesicles (SUVs) prepared from *E. coli* polar lipid extract (EPL) incubated with PspA rods. (B/C) High magnification cryo-EM images of EPL SUVs incubated with PspA rods and EPL SUVs alone, respectively. Fuzzy stretches devoid of a continuous lipid bilayer are marked by arrowheads. (D/E) Scatter plots of EPL vesicle dimensions in the presence of PspA rods and EPL SUVs alone, respectively (grey dashed line represents micrograph field of view). Stacked histogram charts on the right and top axis. (E) EPL forms small uniform vesicles in the absence of PspA, whereas (D) larger deformed vesicles are often observed with a bilayer leaflet separation >5 nm (blue circles). A representative and gallery of diverse EPL vesicles after incubation with PspA rods is shown in **Figure S3**. (F) Statistical analysis of the vesicle perimeters of EPL SUVs control (black) and after incubation with PspA rods (red) and a subset of vesicles with *d* >5 nm (blue) (errors represent SD, n=2524 (black), 2932 (red) and 176 (blue)). (G) Membrane binding of PspA wt (black) and A75S/A78S (red) to negatively charged membranes (40%PG/60%PC) was analyzed by monitoring changes in the Laurdan GP values upon addition of increasing PspA concentrations (error bars represent SE, n=3). The A75S/A78S mutant giving rise to destabilized rods binds with higher apparent affinity to membrane surfaces. See **Figure S5A&B** for the full Laurdan fluorescence spectra. (H/I) Cryo-EM density of PspA+EPL rods superimposed with helical assembly (strand A: red, strand B: cyan) at 7.2 Å resolution. The rods have a width of 275 Å including an inner lumen cylinder density with a thickness of ~40 Å. The inner density is connected to the rod assembly wall by a small tubular stalk that was interpreted as helix α0.

Next, we inspected a large dataset of 1531 cryo-EM images of PspA rod structures in the presence of EPL SUVs in more detail (**Figure S5C**). Based on 2D classification, 65% of rods show a diameter of 275 Å, while only 35% share the narrower diameter at 220 Å observed in the PspA alone rod sample (**Figure S5D&E**). The wider rods (from here: PspA+EPL rods) show a different power spectrum as they form a two-start helix with 13.52 subunits per turn and a pitch of 59.9 Å (**Figure S5F**). Using these helical symmetry parameters, we determined a 7.2 Å cryo-EM density of PspA+EPL rods, which we could fit by the built atomic model of PspA (**Figure 5H, Figure S5G–I**). Notably, PspA+EPL rods enclose a density of lower but significant density forming a separate inner cylinder of ~40 Å thickness. Interestingly, the cryo-EM structure of PspA alone having a smaller diameter also has a pronounced inner density (**Figure S5J&K**), which we attribute to co-purified residual lipids. In addition, we observed a small tubular stalk that protrudes from helix α1 of strand B into the rod lumen (**Figure 5I**), which we interpreted as density for helix α0. Overall, the conformation of PspA determined from PspA+EPL and PspA rods share the same ESCRT-III fold (**Figure S5L&M**). The increased width of PspA+EPL rods are consistent with a lipid bilayer present in the inner lumen and can also be observed in image slices of tomograms of PspA+EPL rods (**Figure S6A**). The experiments with preformed PspA rods and EPL show that the observed vesicle growth is consistent with a membrane remodeling activity and that wider rod structures can potentially accommodate a lipid bilayer in the lumen of the helical rod.

### PspA rods can enclose lipid bilayers and give rise to internalized membrane structures

To study the effect of assembling PspA monomers on liposome membranes rather than preformed PspA rods, we designed a cleavage-inducible assembly variant of PspA (CIA-PspA) that is impaired in rod assembly due to the presence of a N-terminal NusA-tag (**Figure 6A left**). When releasing the NusA-tag from PspA upon 3C protease cleavage, PspA monomers assemble into PspA rods. We found that CIA-PspA remodels liposomes into membrane tubules (**Figure 6A right**) and observed μm-sized clusters of small and large deformed vesicles as well as PspA rods emanating from vesicles (**Figure 6B** and **Figure S6B&C**). In analogy to the incubation with preformed PspA rods, we also observed a pronounced modification in bilayer leaflet separation up to 4.0 nm (**Figure S3D right**). Cryo-tomographic slices (**Figure 6C**) of approx. 50 % of CIA-PspA rods show clear tubular lipid bilayer density protruding from the vesicle into the lumen of the rods, while other smaller diameter CIA-PspA rods did not enclose tubular lipid bilayers (**Figure S6D**). Interestingly, the rods were always attached to small vesicles with a consistent polarity with respect to the vesicles. By superimposing the PspA+EPL rods with CIA-PspA rods in the tomographic slices, we infer that the rods attach to the vesicles with their negatively charged end **(Figure 6C** and **Figure S5N**). In conclusion, PspA rods can incorporate a lipid bilayer into the central lumen producing constricted positively curved membrane tubules.

**Figure 6:**
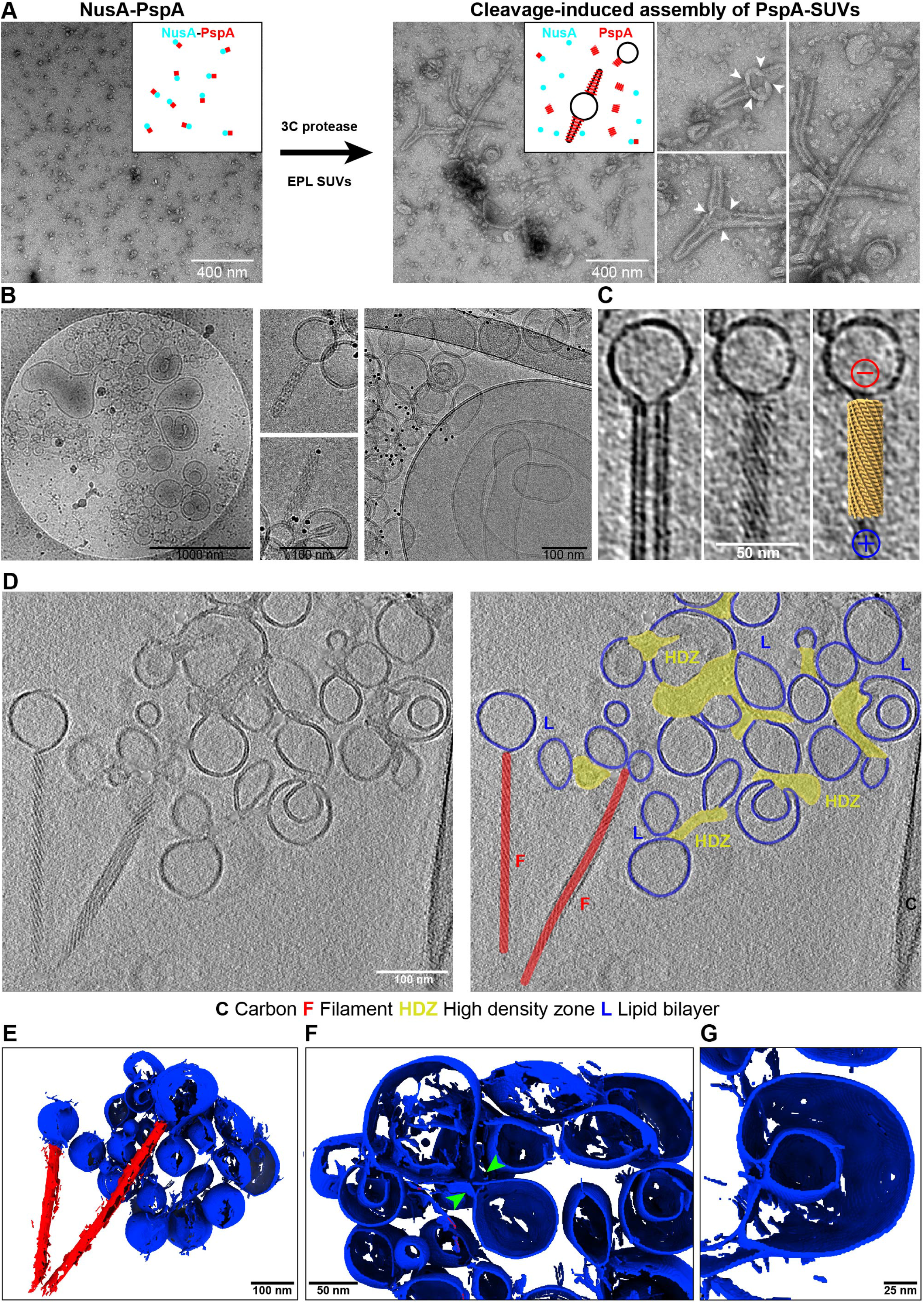
Structures of membrane-attached PspA rods formed by cleavage-induced assembly (CIA-PspA). (A) Negatively stained electron micrographs of NusA-PspA and CIA-PspA. After 3C protease addition to NusA-PspA (globular structures) in the presence of EPL SUVs, PspA spontaneously forms membrane tubules attached to vesicles (white arrowheads point to tubule-vesicle contact sites). (B) Cryo-EM micrographs of CIA-PspA in presence of EPL SUVs. Left: Overview image indicates structural diversity of vesicles in this sample (see also **Figure S6**). Center and right: Detailed images of rods attached to vesicles and fused or deformed vesicles. (C) Tomogram slices of a CIA-PspA rod attached to a vesicle. Left: central slice showing a lipid bilayer in the inner lumen. Center: Top slice showing striations of the outer helical lattice. Right: Top slice superimposed with a model of the polar PspA+EPL rod. The negatively charged end of the rod faces the vesicle. (D) Left: representative cryo-electron tomogram of CIA-PspA and a cluster of EPL SUVs (**Video S1**). Right: the same tomogram with superimposed 2D segmentation showing rods in red, high-density zones (HDZ) in yellow and lipid bilayers in blue. (E) 3D segmentation of the tomogram shown in (D). The vesicles and HDZs form an interconnected 3D lipid/PspA network. (F) Zoomed view of the interconnected lipid/PspA network with two fusing vesicles (green arrow heads) in the center. (G) Detailed view of a larger vesicle with an enclosed budding vesicle.

However, mixtures of PspA and EPL SUVs have a broad distribution of vesicle sizes and membrane structures. In-depth analysis of a representative tomogram by continuous z-slices reveals that multiple vesicles are often connected, thus forming an intricate system of membranous cisternae linked together by proteinaceous high-density zones (HDZ), where distinct lipid bilayer density is absent (**Figure 6D, Video S1**) giving rise to the observed fuzzy stretches in the 2D images described above. Nevertheless, HDZs can often be found close to areas of high lipid curvature at the cone-shaped and tubule-like ends of pear-shaped vesicles (**Figure 6E&F**). Close to HDZs, often internalized vesicles can be found connected to a larger vesicle suggesting an internal budding process (**Figure 6G**). Together with the observed structural ESCRT-III homology, co-reconstitution with lipids demonstrates that PspA is capable of mediating vesicle growth as well as membrane constriction and fission in dependance on the initial oligomeric state, and can, therefore, be considered a *bona fide* member of the ESCRT-III family of membrane remodeling proteins.

## Discussion

The propensity of PspA to form higher-order structures has been shown early on (Hankamer et al., 2004; Kobayashi et al., 2007), yet how oligomerizations contributes to the basic physiological PspA function of maintaining membrane integrity remained uncertain. Here, we determined the 3.6 Å resolution cryo-EM structure of PspA rod assemblies and systematically studied the interaction of PspA with membranes. Due to similar structural and functional properties of PspA and eukaryotic ESCRT proteins, we here identified PspA as a bacterial member of the evolutionary conserved ESCRT-III superfamily of membrane remodeling proteins.

While EcoPspA forms rings rather than rods (Hankamer et al., 2004; Kobayashi et al., 2007), the physiological importance of ring *vs*. rod structures remains an open question, as these structures have not been observed in their native context, yet. Structurally, rods and rings are closely related, and conversion can be easily envisioned by accommodating small angular changes in subunit-subunit interactions. Using *in vivo* fluorescence microscopy, IM30 and PspA were shown to form dynamic localized *puncta* in *Synechocystis* and *E. coli* (Engl et al., 2009; Gutu et al., 2018; Jovanovic et al., 2014; Yamaguchi et al., 2013), which is consistent with formation of rings, transient rod or other higher-order structures associated with cellular membranes (Junglas and Schneider, 2018; Siebenaller et al., 2019). We here, and others, observed that such PspA assemblies were preferentially found at the cell poles, most likely to protect vulnerable cell membrane regions from dissipation of the proton motive force (Engl et al., 2009).

Due to revelation of PspA as a member of an ESCRT-III superfamily, we wondered whether there are more structural similarities beyond the identified homology and fold. Purified ESCRT-III proteins are known to assemble into a variety of large EM-observable structures, amongst them strings, rings, coils, sheets, filaments and domes (Bodon et al., 2011; Ghazi-Tabatabai et al., 2008; Lata et al., 2008). Due to the large number of isoforms of eukaryotic ESCRT-III proteins, the complexity of assembling hetero-complexes is significantly increased, and multiple combinations have been investigated (Henne et al., 2011; Huber et al., 2020; Pfitzner et al., 2020). A structural motif of striking similarity found in PspA as well as in eukaryotic ESCRT structures is the perpendicular packing of helix α5 against the α1/α2 hairpin (**Figure 4**). Additionally, helix α4 establishes important intersubunit contacts in the central core of the polymers, which is reflected in a stretch of residues with properties conserved between PspA/IM30 and ESCRT III proteins (**Figure 3**). While the AAA-ATPases PspF and Vps4 disassemble *Eco*PspA or eukaryotic ESCRTs, respectively, PspF is not encoded in *Synechocystis* and other PspA-interacting ATPases have not be identified. Yet, a currently enigmatic ATPase may be involved in regulating the PspA activity in *Synechocystis* given the central importance of Vps4 in the ESCRT-III membrane remodeling pathway.

Despite the matching structural organization of PspA and ESCRT-III proteins, we further assessed whether PspA and ESCRT-III proteins have functional similarities. Consistent with the function of ESCRT-III being involved in membrane remodeling processes, PspA co-incubated with EPL revealed two noteworthy details: PspA binding to membrane surfaces induced a leaflet separation >5 nm and the appearance of remodeled liposomes. Observations include PspA-mediated vesicle growth as well as formation of membrane tubules and internalized membrane structures resulting from fusion, constriction and fission events. The widening of the lipid bilayer likely is a consequence of PspA rod disassembly and the formation of a PspA carpet on the membrane surface, as recently described for IM30 (Junglas et al., 2020). Membrane adhesion and partial incorporation of PspA molecules into the membrane will lead to the here observed thickening of the lipid bilayer. The observed bilayer structures could represent a protected form of the lipid bilayer minimizing, for instance, leakage of protons under imposed membrane stress conditions (Junglas et al., 2020; Kobayashi et al., 2007). Electrostatic surface analysis of the PspA rod structure reveals an acidic and a basic end, which could provide a rationale for PspA’s observed preference of binding to negatively charged membrane surfaces (Kobayashi et al., 2007; McDonald et al., 2015, 2017). Yet, our membrane interaction studies show that PspA is also capable of binding to net uncharged membrane surfaces, and, therefore, membrane interaction of PspA appears to be less charge-dependent than observed for *Syn*IM30 (Hennig et al., 2015).

The imaged liposomes also show high-density zones devoid of any visible organized lipid bilayer structure, often with PspA rods attached to the vesicle surface. The corresponding regions contain a grainy texture suggestive of floating PspA molecules dissociated from the rod into smaller units. As they make up the linkages of a continuous vesicular lipid network, they likely constitute lipid transfer zones (LTZ), enabling lipid movement between vesicles. In this role, LTZs may locally replace the bilayer structure while giving up the characteristic phospholipid barrier structure. In many aspects, the here observed LTZs and PspA’s membrane activity is reminiscent of the detergent-like membrane interaction of linear amphipathic antimicrobial peptides (reviewed in (Bechinger and Lohner, 2006)). In previous experiments, it has been observed that interaction of *Syn*IM30 with membranes disturbs the bilayer structure and allows small molecules to enter liposomes (Hennig et al., 2015) and an antimicrobial peptide-like activity has been discussed (Thurotte and Schneider, 2019). Importantly, a detergent-like behavior of PspA does not exclude formation of stabilizing membrane carpets and formation of destabilizing pore-like structures within a membrane.

The here determined cryo-EM structures of PspA and PspA+EPL rods bear most apparent similarity with the determined IST1/CHMP1B ESCRT-III assembly (McCullough et al., 2015). More recently, IST1/CHMP1B could also be reconstituted with negatively charged lipids that were found in the lumen of the assembly (Nguyen et al. 2020). In presence of lipids, the diameter of the IST1/CHMP1B tubular assemblies increased from 240 to 280 Å leading to the hypothesis of high and low constriction states, respectively. In analogy to the IST1/CHMP1B as well as *Chlamydomonas* IM30/Vipp1 rods (Theis et al., 2019), we here observed that PspA rods can include lipid bilayers in the lumen, accompanied by changes in diameter from 220 to 275 Å, which therefore, represent different constriction states. Noteworthy, our preparations of PspA alone also revealed a significant variation of the diameters of the rods. This structural flexibility may be an important property of PspA to accommodate different bilayer intermediate configurations during the lipid interaction process. Furthermore, in the presence of lipids we detected density for helix α0 in the PspA rod structure consistent with folding of N-terminal helix peptides of PspA and IM30 in presence of lipid bilayers (McDonald et al., 2015, 2017). Taken together, PspA rods are strong candidates for membrane remodeling proteins generating positive curvature, as IST1/CHMP1B and PspA assemblies share the same principal rod architecture, lipid topology and changes in diameters.

In order to describe the biogenesis of the observed membrane fusion, constriction and fission activities, we sorted the set of recorded membrane events to enable interpretation in a sequential manner. Initially, PspA assembles in rods free in solution or starting from a membrane anchored platform (**Figure 7 top**). The PspA+EPL intermediate resolution cryo-EM structure and cryo-tomograms show that these membrane-attached rods incorporate lipid bilayer tubules into the observed wider structures. The polar rods grow at the positively charged ends at the tip until they may encounter other free vesicles containing negatively charged lipids. In this contact area, ordered PspA rods start to disassemble at their positive end and initiate formation of LTZs, connecting two vesicular structures (**Figure 7 center**). After this docking event, the vesicular lobes form the structure resembling an hour-glass linked by the PspA LTZ. This hour-glass structure likely represents a critical switch structure that we propose can undergo two principal pathways: In the first case, lipid movement is allowed over the LTZ and the donor membrane of vesicle 1 is in close spatial proximity to the acceptor membrane of vesicle 2. Then, lipid transfer across a newly formed continuous lipid structure can take place thereby fusing the bilayers of vesicle 1 and 2. Upon relaxation of the lipid bilayer structure, larger vesicles consisting of the fused membranes of vesicle 1 and 2 have formed (**Figure 7 bottom left**). In the second case, lipid movement is allowed over the LTZ and the donor membrane of vesicle 1 is not in closest spatial proximity to the membrane of vesicle 2. Here, the membranes of vesicle 1 and 2 do not form a continuous bilayer, and as a result, the membrane of vesicle 1 buds into the lumen of vesicle 2, resulting in two separate membrane systems. In consequence, a cross-membrane transfer of vesicle 1 into the lumen of vesicle 2 and membrane fission is observed (**Figure 7 bottom right**), reminiscent of ESCRT-III inward vesicle budding events. We would like to point to out that in our *in vitro* reconstituted samples, we classified these two principal cases although we also observed multiple intermediate events and combinations of these two outlined pathways, *e.g.* including a symmetric cross-membrane vesicle transfer and multiple others (**Figure S7**). According to this fusion-fission switch model, the critical step that determines the fate of fusion or fission may be the spatial proximity of the donor and acceptor membrane that enables or disables the formation of a continuous lipid bilayer. In both of the described activities, PspA’s mode of action results from the ability to move lipids across the LTZ.

**Figure 7:**
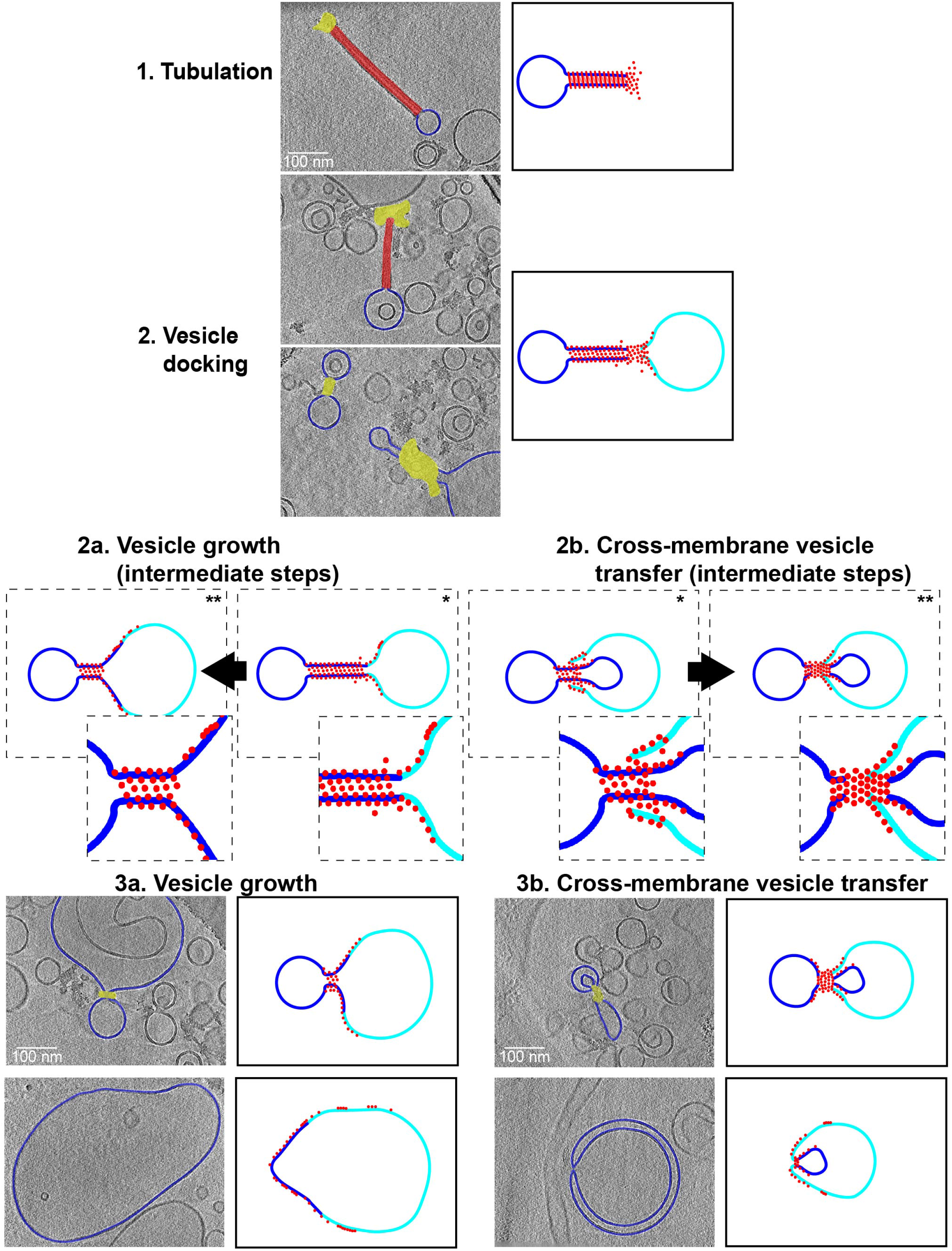
Model of PspA membrane remodeling activity resulting in vesicle growth and cross-membrane vesicle transfer. A potential sequence of events leading to vesicle growth and cross-membrane vesicle transfer supported by cryo-EM images. Left column: cryo-electron tomograms of PspA interacting with vesicles superimposed with 2D segmentation of the region of interest (rods in red, lipid transfer zones in yellow and lipid bilayers in blue) (Corresponding non-segmented tomogram slices shown in **Figure S7**). Right column: Simplified model (blue: donor membrane, cyan: acceptor membrane, red: PspA). The initial two steps are identical for vesicle growth and cross-membrane vesicle transfer: 1. Tubulation, PspA assembles into rod structures anchored with one vesicle in a polar manner. 2. Vesicle docking, a second vesicle will be contacted by the PspA rod linking the two vesicles. Switch point leading either to 2a. or 2b. that represent proposed intermediate steps (dashed box with zoomed insets of lipid transfer zones): Depending on the spatial proximity of the donor and acceptor membrane within the lipid transfer zone 3a. vesicle growth or 3b. cross-membrane vesicle transfer can occur. Relaxation of vesicular constriction leads to a large fused vesicle. Donation of lipid vesicle into lumen of juxtaposed vesicle leads to an internalized membrane structure.

The here described fusion-fission switch model explains a multitude of the observed membrane activities of PspA. Upon occurrence of membrane leakage, PspA-induced vesicle growth can potentially patch disturbed membranes and restore full membrane integrity thereby protecting bacterial membranes, as it has been hypothesized previously for the bacterial dynamin like protein DynA after treatment of membranes with pore-forming agents (Guo and Bramkamp, 2019). In addition to this active repair mechanism, PspA likely is also capable of passive membrane protection by forming a protective carpet on membranes, as described above. Furthermore, PspA-induced cross-membrane vesicle transfer can result in the formation of intricate internalized membrane systems reminiscent of thylakoid membrane stacks that is known to be mediated by the PspA homolog IM30 (Heidrich et al., 2017; Vothknecht et al., 2012). Similarly, inward vesicle budding as it has been attributed to eukaryotic ESCRT-III action during multivesicular body biogenesis, could also be envisioned by the process of cross-membrane vesicle transfer. In case PspA’s mode of action was transferable to eukaryotic ESCRT-III proteins, the determined structure of IST1/CHMP1B polymer would be directly compatible, as it has also been shown to bind lipids in the lumen of the assembly. In contrast to lipid reconstitutions mediated by several eukaryotic ESCRT proteins, the homo-oligomer of PspA alone induces large-scale changes to liposome morphology, including growth, constriction and cross-membrane vesicle transfer. Thereby, it represents a minimal bacterial membrane remodeling system that is capable of triggering membrane fusion and fission *in vitro*.

### Limitations of the study

While this study establishes that PspA is a member of the ESCRT-III family, this membership has now been confirmed for PspA related bacterial IM30/Vipp1 proteins by two independent studies: first, derived from bioinformatic analysis and followed by single-particle cryo-EM of IM30/Vipp1 rings revealed a conserved ESCRT-III fold (Liu et al., 2020). Second, the cryo-EM single-particle structure of IM30/Vipp1 rings as well as *in situ* cryo-EM studies revealed the presence of Vipp1 rods in *Chlamydomonas reinhardtii* (Gupta et al., 2020). Our study adds functional details on the membrane remodeling activities to the obtained structures, however, it also raises a series of new questions:

1. Are the observed PspA’s remodeling activities fully transferable to the diversity of activities of eukaryotic ESCRT-III proteins?
2. Do, in analogy to eukaryotic ESCRT-III, regulators exist *in vivo* that direct PspA’s membrane remodeling activities to fulfill different functions in membrane organization, repair and maintenance?
3. Do prokaryotic IM30 or LiaH share the same mechanisms of membrane remodeling?

To address these open questions, it will be important to identify further mechanistic details of structural intermediates of the PspA membrane remodeling system responsible for maintaining bacterial membrane integrity *in vitro* and *in vivo*.

## Acknowledgments

We acknowledge support with microscope operation, image acquisition and preparation of Figure 6 by Julio Ortiz in addition to helpful discussions. The authors gratefully acknowledge the computing time granted through JARA on the supercomputer JURECA at Forschungszentrum Jülich (Krause, 2019). This work was funded by the Max-Planck Graduate Center at the Max Planck institutes and the University of Mainz. This work has been initially supported by iNEXT, grant number 653706, funded by the Horizon 2020 program of the European Union.

## Author contributions

B.J., D.S. and C.S. designed research. B.J. and L.S. purified the proteins. B.J., T.H. and M.C. prepared cryo-EM samples. T.H., M.C. and B.J. operated the electron microscopes. B.J. and S.T.H. determined the cryo-EM structures. D.M. and B.J. built the refined atomic model. R.H. performed mutation experiments. B.J. and N.H. performed membrane-binding experiments. L.S. conducted the *in vivo* experiments and fluorescence microscopy. B.J., D.S., and C.S. wrote the manuscript with input from all authors.

## STAR Methods

Detailed methods are provided in the online of this paper and include the following:

- Key Resource Table
- Resource Availability

- Lead Contact
- Materials availability
- Data and code availability
- Experimental Model and Subject Details

- Bacterial cells
- Methods Details

- Expression and purification of PspA
- Lipid reconstitution
- Membrane binding assay
- Cleavage-induced assembly of PspA rods (CIA-PspA)
- Fluorescence microscopy
- Negative-staining electron microscopy
- Electron cryo-microscopy
- Electron cryo-tomography (cryo-ET)
- Image processing and helical reconstruction
- Cryo-EM map interpretation and model building
- Cryo-ET image processing
- Image analysis of membranes
- Quantification and Statistical Analysis

## Supplemental Videos

**Video S1. Cryo-electron tomogram of CIA-PspA interacting with EPL vesicles (related to Figure 6D–G).**

## Supplemental Figures

**Figure S1.**
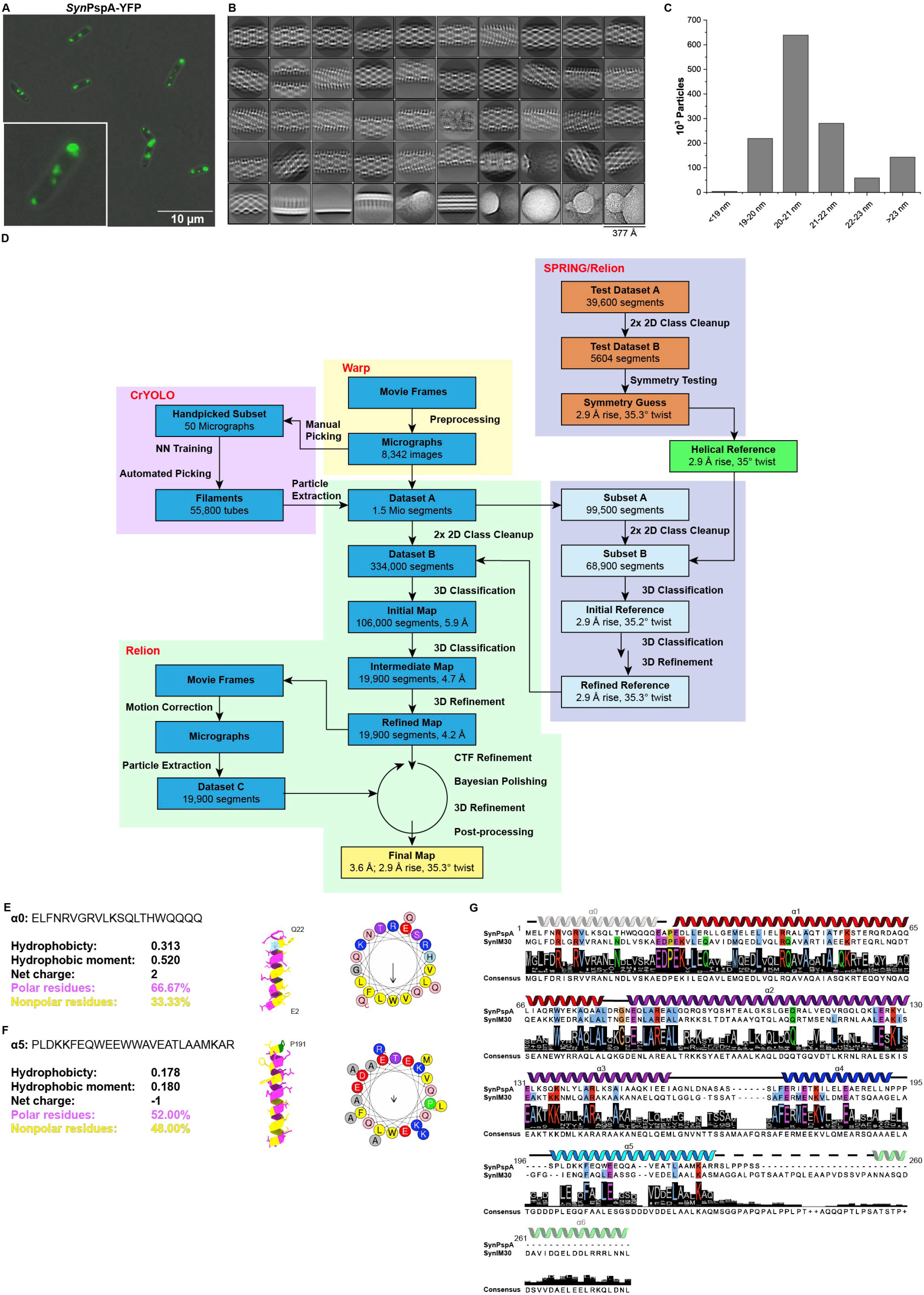
Fluorescence microscopy images, processing details and sequence alignment of PspA (related to Figure 1 – 3). (A) Epifluorescence images of *E. coli* Δ*pspA* cells expressing *Syn*PspA YFP-tagged at the C-terminus. Low-level expression of the fusion proteins was induced after addition of 0.02% L-arabinose. Green fluorescent foci are visible within cells, representing higher-ordered PspA assemblies. Typically, *E. coli* cells contain two foci (71%, of n=197 cells), one localized at each cell pole. In case of dividing cells, fluorescent foci are also visible at the division septum (*i.e.* the new cell pole) within each daughter cell. (B) Gallery of 2D class averages from initial 1.5 million segment data set. (C) Histogram showing distribution of measured widths of PspA rods. (D) Basic image processing workflow of PspA structure determination: PspA rods were segmented and a subset subjected to symmetry determination in SPRING followed by symmetry refinement in RELION. Once the symmetry was determined, RELION was used for final 3D structure determination of PspA including CTF refinement and Bayesian polishing. (E) Helix wheel projection of the predicted helix α0. (F) Helix wheel projection of helix α5. Projections were generated with HELIQUEST (Gautier et al., 2008). (G) A conservation query for the PspA/IM30 protein was subjected to the ConSurf server (Ashkenazy et al., 2016) using the *Syn*PspA and the *Syn*IM30 sequence as input. The output of 250 sequences was aligned using ClustalW (Larkin et al., 2007) The resulting consensus sequence together with the *Syn*PspA and *Syn*IM30 were visualized in Jalview (Waterhouse et al., 2009) and superimposed with the PspA/IM30 secondary structure using the ESCRT-III helix nomenclature. Residues with >90% identity are colored by their properties according to the ClustalX color scheme.

**Figure S2.**
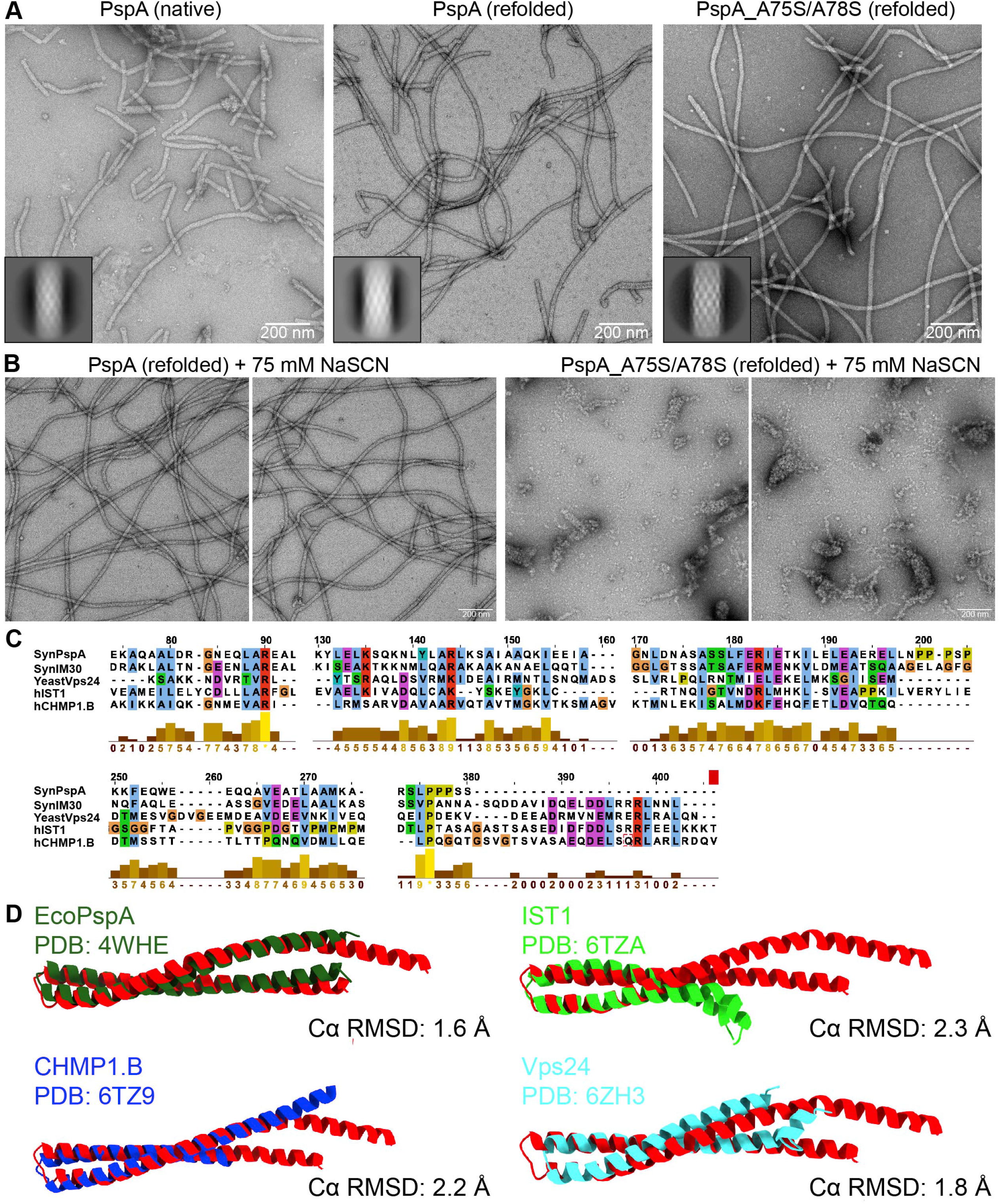
Negative-staining electron micrographs of PspA and comparison of α1/α2 hairpin motif of PspA with eukaryotic ESCRT-III structures (related to Figure 2&4). (A) Micrographs of negatively stained samples of PspA after native extraction, and PspA as well as PspA_A75S/A78S after purification under denaturing conditions and subsequent refolding. The inset shows a representative 2D class average of each sample. (B) Micrographs of negatively stained samples of PspA and PspA_A75S/A78S after treatment with 75 mM NaSCN. After overnight dialysis against sodium phosphate buffer containing 75 mM NaSCN, rod formation is not affected in case of wt PspA, while formation of PspA_A75S/A78S rods is impaired. (C) Sequence alignment of PspA/IM30 with eukaryotic ESCRT-III proteins with bar graphs showing the alignment quality score as a measure of sequence similarity. Sequences were aligned using the T-Coffee webserver (Di Tommaso et al., 2011) and visualized with JalView (Waterhouse et al., 2009). (D) Structural superposition of the PspA hairpin with *Eco*PspA, IST1, CHMP1-B and Vps24. CA RMSD of residues approx. 50 – 90 (central hairpin) was calculated in Chimera using the “Match**→**Align” command.

**Figure S3.**
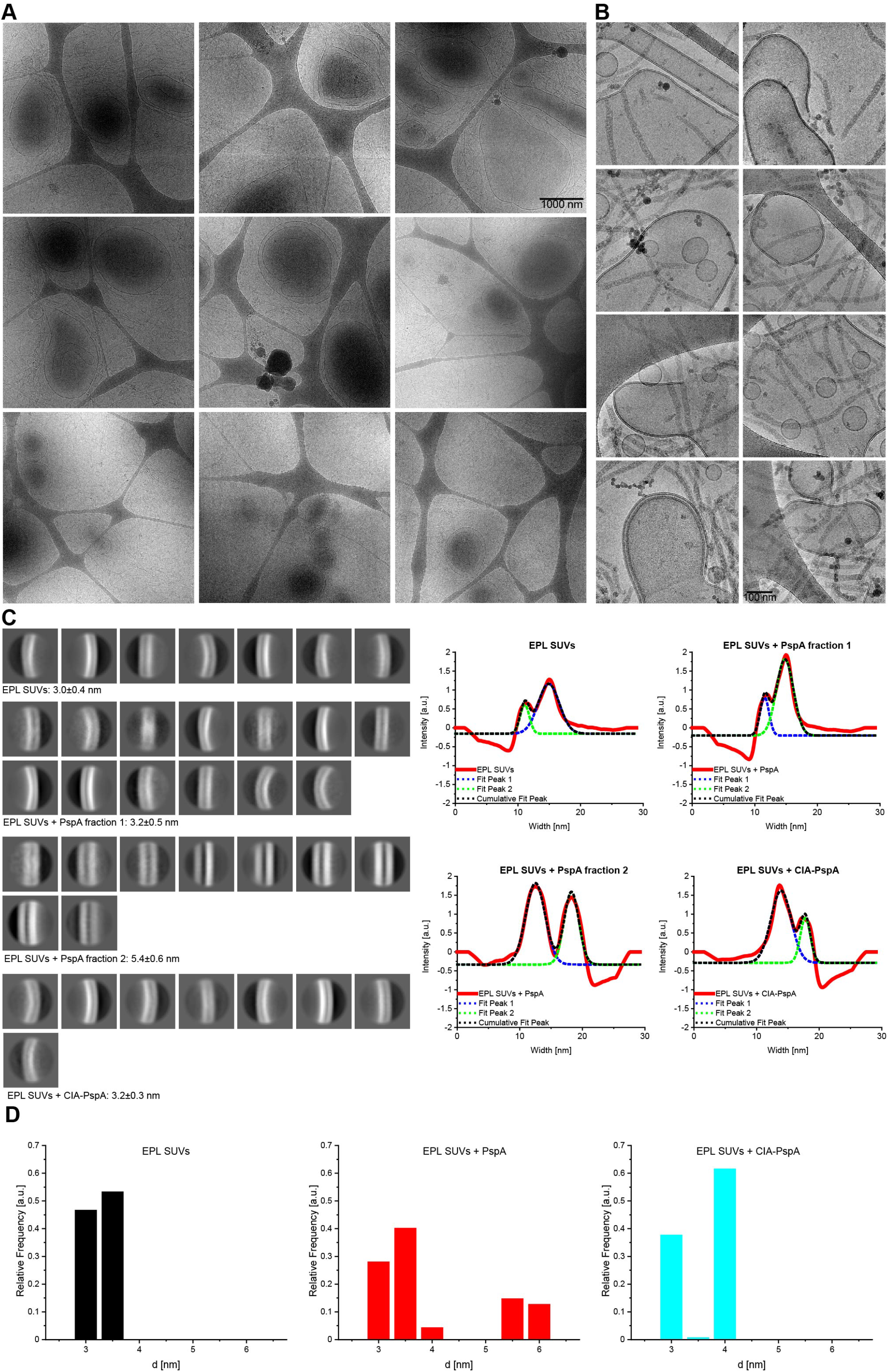
Gallery of cryo-micrographs showing *E. coli* polar lipids (EPL) small lamellar vesicles (SUVs) after incubation with PspA rods and image analysis (related to Figure 5). (A) Electron cryo-micrographs of EPL SUVs in the presence of PspA rods at low magnification. (B) EPL SUVs in the presence of PspA rods at high magnification. (C) Left. Class averages of bilayer segments of EPL, EPL+PspA and EPL+CIA. The average bilayer leaflet separation *d* denotes the average over all class averages per group with SD (n=number of respective class averages). Right. Representative density profiles of the respective class averages including the fitted peaks that were used to determine the bilayer leaflet separation. (D) Relative frequency of the determined bilayer leaflet separation based on the number of bilayer segments per class binned to 0.5 nm groups.

**Figure S4.**
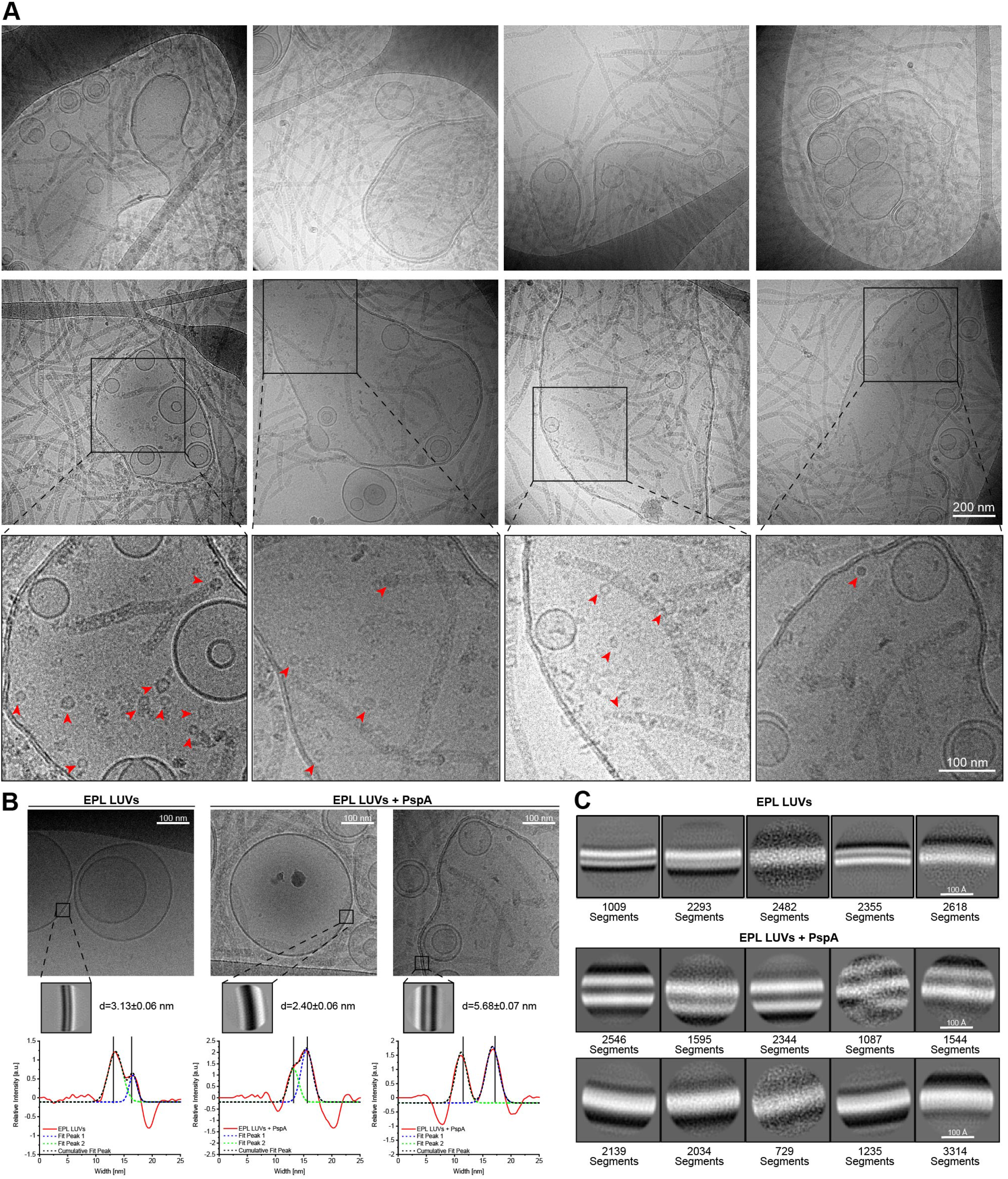
Gallery of cryo-micrographs showing *E. coli* polar lipid (EPL) large unilamellar vesicles (LUVs) after incubation with PspA rods and image analysis (related to Figure 5). (A) Electron cryo-micrographs of EPL LUVs in the presence of PspA rods at low magnification. Bottom row: magnified insets of the last row of LUV images. Red arrows indicate smaller oligomeric structures such as rings. (B) Top. In the absence of PspA, EPL large unilamellar vesicles (LUVs) are uniform and round, whereas larger and deformed vesicles were observed in presence of PspA showing a distinct bilayer leaflet separation in addition to a fraction of unaltered uniform vesicles. Bottom. Detailed analysis of bilayer leaflet separation *d* in EPL vesicles determined from density profiles of class averages from segmented bilayers. Only class averages with clear features were used for the measurement. (n=3 class averages, errors represent SD). (C) Class averages in absence and presence of PspA.

**Figure S5.**
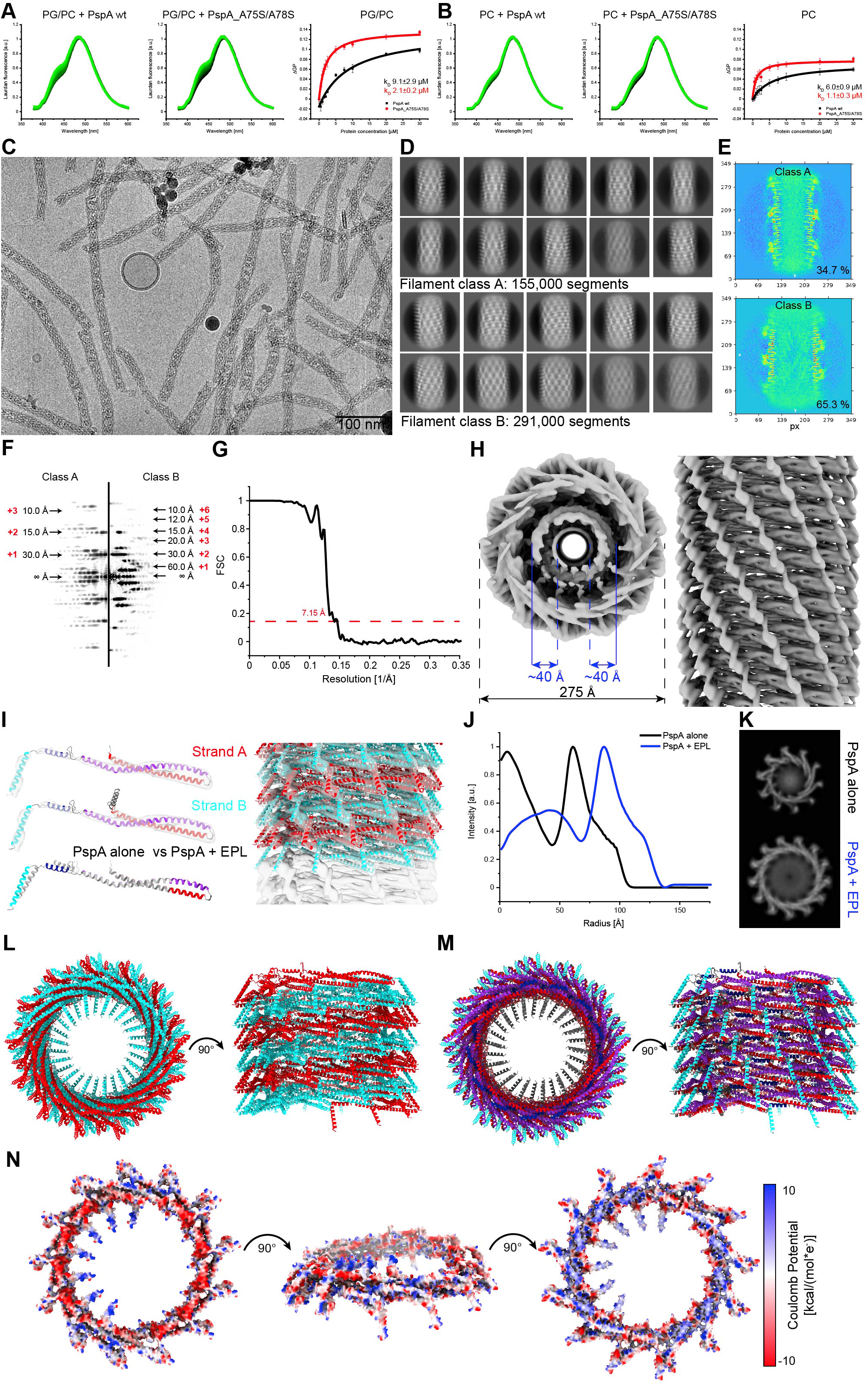
Membrane binding of PspA measured by fluorescence spectroscopy and cryo-EM structure of PspA rods in presence of EPL (related to Figure 5). (A/B) Laurdan fluorescence spectra for lipid binding of PspA wt and A75S/A78S to negatively charged DOPC/DOPG (60/40) (A) or net uncharged DOPC membranes (B). Spectra with increasing protein concentrations from 0 - 30 μM are shown (color coded with dark green to light green). Binding of PspA wt and mutant leads to an apparent increase in lipid order, indicated by the relative increase of the emission band around 440 nm. The binding curves constructed from the corresponding GP values show that binding of both variants is largely independent of the net membrane charge. Yet, both proteins have a slightly increased propensity to bind to negatively charged membranes, and the mutant, where the rods are destabilized, binds with higher apparent affinity to membrane surfaces. An increase in lipid order induced by protein binding cannot be detected if the membrane is already largely in the ordered phase, thus this assay cannot be used to detect PspA binding to EPL liposomes. Error bars in the binding curves (right column) represent SE (n=3). (C) Representative electron cryo-micrograph of PspA rods in the presence of EPL small unilamellar vesicles (SUVs). (D) Class averages of rods grouped in two classes. (E) y-z slice of the 3D reconstruction of each rod class. The percent value in the bottom-right location indicates the segment ratio contributing to the respective 3D reconstruction. (F) Power spectra of 2D class averages of the two rod classes (class A left, class B right) indicating layer lines up to 10 Å resolution. Red numbers represent the predicted Bessel order of each layer line. (G) Fourier shell correlation of rod class B indicating a 7.2 Å resolution at the 0.143 cutoff. (H) Top and side view of cryo-EM density of rod class B with measurement of the width and thickness of the inner density. (I) Left: segmented cryo-EM density superimposed on built model of PspA (strand A:22-217, strand B: 1-217) with colored α-helices in ribbon presentation (α1: red, α2/α3: purple, α4: blue, α5: cyan) and comparison of the model of PspA alone *vs.* PspA+EPL. Right: density of PspA+EPL rods with fitted helical assembly in side view. The assembly is a two-start helix with 4.4 Å rise and 26.6° twist (strand A in red, strand B in cyan). (J) Radial density profiles of PspA alone and PspA+EPL (class B) rods. (K) Corresponding top view projection of PspA alone and PspA+EPL (class B) rods that were used to determine radial density profiles. (L) Helical assembly of PspA+EPL in top and side view (strand A in red, strand B in cyan). (M) Helical assembly of the PspA+EPL rod in side and top view with colored α-helices in ribbon presentation (α1: red, α2/α3: purple, α4: blue, α5: cyan). (N) Electrostatic surface potential of PspA+EPL rods. The assembly consists of two unique ends with basic and acidic surfaces and pronounced positively charged spots in the inner volume.

**Figure S6.**
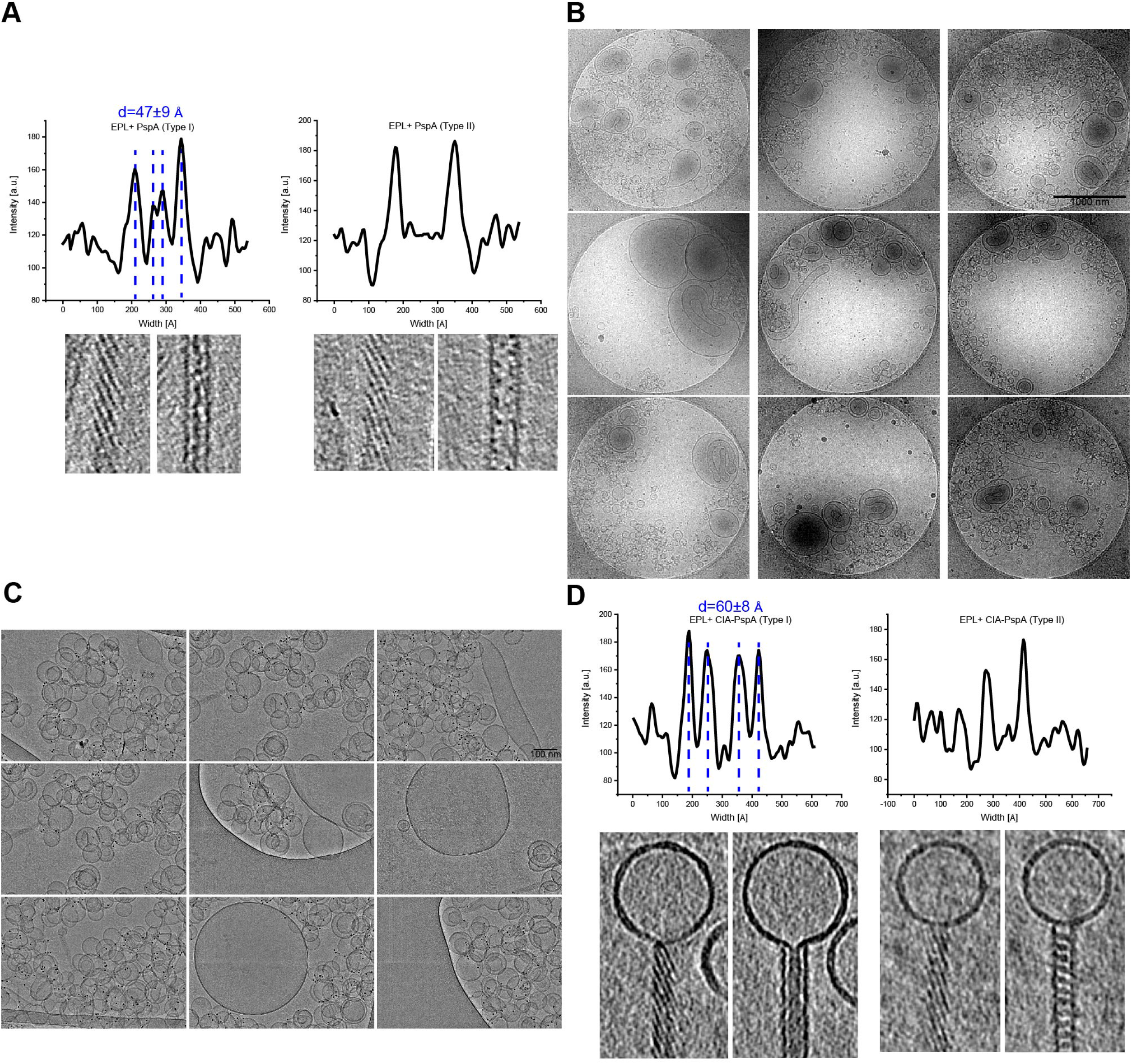
Density profiles of PspA rod tomogram slices and gallery of EPL SUVs after incubation with CIA-PspA (related to Figure 6). Top: representative density profiles of PspA rods in the presence of EPL SUVs. Type I rod profiles show a distinct double-U pattern with peak-to-peak distances of 47±9 Å (errors represent SD, n=6), while the profiles of type II rods show a single-U pattern. Bottom: top and central tomogram slices of the respective rods. (B) Electron cryo-micrographs of EPL SUVs in the presence of CIA-PspA at low magnification. (C) EPL SUVs in the presence of CIA-PspA rods at high magnification. (D) Top: representative density profiles of CIA-PspA rods in the presence of EPL SUVs. Type I rod profiles show a double-U pattern with peak-to-peak distances of 60±8 Å (errors represent SD, n=8), while the profiles of type II rods show a single-U pattern. Bottom: top and central tomogram slices of the respective rods. The density of the observed double-U pattern is compatible with a lipid bilayer present in the inner lumen of the PspA rod.

**Figure S7.**
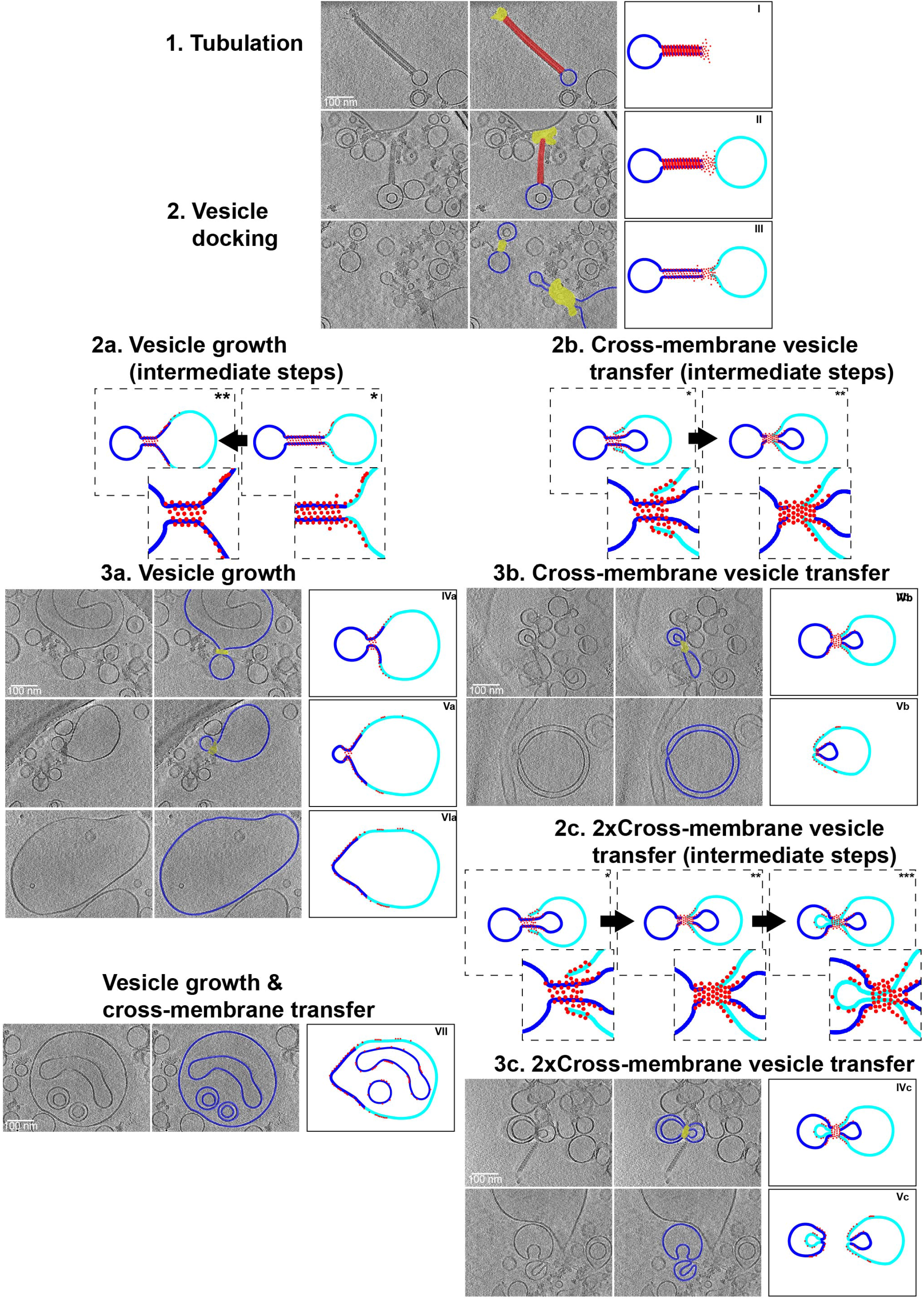
Detailed sequence of observed membrane topologies leading to a model of vesicle growth and cross-membrane vesicle transfer (related to Figure 7). A potential model of events leading to vesicle growth and cross-membrane vesicle transfer supported by cryo-EM images (Extended view of Figure 7). Left column: image slices of cryo-electron tomograms of PspA interacting with vesicles. Middle column: corresponding slices superimposed with 2D segmentation of the region of interest (rods in red, high-density zones in yellow and lipid bilayers in blue). Right column: simplified model (blue: donor membrane, cyan: acceptor membrane, red: PspA). For details see the main text.

